# Conserved intrinsically disordered region of DNAJB6 dictates its surveillance of FG-Nup condensates

**DOI:** 10.1101/2025.10.20.683411

**Authors:** Tessa Bergsma, Maiara Kolbe Musskopf, Alejandro Feito, Paola Gallardo, Mathieu E. Rebeaud, E. F. Elsiena Kuiper, Nuria H. Espejo, Andres R. Tejedor, Jarmo Feenstra, Sidath M. Y. Fernando, Anton Steen, Rifka Vlijm, Jorge R. Espinosa, Harm H. Kampinga, Liesbeth M. Veenhoff

**Affiliations:** European Research Institute for the Biology of Ageing, University of Groningen, University Medical Center Groningen, 9713 AV, Groningen, the Netherlands; Biomedical Sciences, University of Groningen, University Medical Center Groningen, 9713 AV, Groningen, the Netherlands; University Complutense of Madrid, Physical Chemistry Department, Avenida Complutense s/n 28040, Madrid, Spain; Multidisciplinary Institute, Complutense University of Madrid, Paseo Juan XXIII, 1, Madrid 28040, Spain; Centro Andaluz de Biología del Desarrollo, Universidad Pablo de Olavide, Avenida Rectora Rosario Valpuesta 1, 41089 Dos Hermanas, Sevilla, Spain; Institute of Physics, School of Basic Sciences, École Polytechnique Fédérale de Lausanne, CH-1015, Lausanne, Switzerland; Mechanisms of Cellular Quality Control, Max Planck Institute of Biophysics, Frankfurt am Main, Germany; Molecular Biophysics, Zernike Institute for Advanced Materials, University of Groningen, 9747 AG Groningen, the Netherlands; Yusuf Hamied Department of Chemistry, University of Cambridge, Lensfield Road, Cambridge CB2 1EW, United Kingdom; Phasica Biosciences S.L., Calle Velazquez, 27, 28001 Madrid, Spain

**Author notes:** Equal contribution (Tessa Bergsma, Maiara Kolbe Musskopf).

**Keywords:** DNAJB6, FG-Nups, phase transition, intrinsically disordered proteins, aggregation

## Abstract

Molecular chaperones are known for their role in preventing protein aggregation and assisting proteins in reaching their structurally functional state. DNAJB6, a J-domain protein that partners with Hsp70s and nucleotide exchange factors, is very potent in preventing amyloid formation of proteins with large intrinsically disordered regions (IDRs), including several disease-associated proteins. Complementary to this, we recently demonstrated a role for DNAJB6 in surveilling FG-Nucleoporins (FG-Nups) phase transitions and highlighted its role in nuclear pore complex assembly. We expand on this by showing that this activity of phase state surveillance is directed to several FG-Nups and shared with the closely related DNAJB2 and DNAJB8. We demonstrate that the surveillance mechanism of DNAJB6 is encoded in an unusually highly conserved IDR that promotes the formation of stable, gel-like assemblies of the chaperone itself. These assemblies likely provide a stable environment that can outcompete stable homotypic FG-Nup interactions and instead favors multivalent heterotypic chaperone:FG-Nup interactions. The evolutionary conservation of the DNAJB6-IDR, mutant analyses from both experimental *in vitro* and in cell data, and multiscale molecular dynamics simulations suggest that the sequence space for encoding stable gel-like assemblies is narrow and optimized to avoid self-aggregation while providing potent anti-amyloidogenic capacity.

## Introduction

Intrinsically disordered proteins (IDPs) are present in all kingdoms of life, and they have gained significant attention due to their unique properties and functional capabilities^1,2^. Unlike traditional proteins that adopt stable three-dimensional structures, IDPs lack a stable structure under physiological conditions^3^. This inherent flexibility allows them to partake in dynamic multivalent interactions forming homotypic or heterotypic biomolecular condensates within the cellular environment. These condensates facilitate crucial biological processes such as regulating gene expression, compartmentalizing metabolic reactions and transiently and locally concentrating molecules^4,5^.

IDPs can adopt ensembles of rapidly interconverting structures. This behavior is dependent on the IDP sequence and is also highly context-dependent, where the context is defined by, amongst others, the concentration of the IDP, macromolecular crowding, temperature, binding partners, pH, and post-translational modifications^4^. The self-assembly of IDPs into dynamic condensates can occur through, amongst other principles, liquid-liquid phase separation (LLPS), in which a homogeneous solution of molecules demixes into distinct, coexisting liquid-like phases. Not uncommonly, liquid-like condensates transition into a gel-like phase, in which the interactions become less dynamic^6–9^. In fact, structural motifs termed Low-complexity Aromatic-Rich Kinked Segments (LARKS), which are commonly found in hydrogel-forming IDPs, promote phase transition into reversible semisolid hydrogels that organize in a cross-ꞵ pattern, but the paired kinked ꞵ-sheets of their LARKS are less strongly bound than the paired ꞵ-sheets found in amyloid fibrils^10^. However, under certain conditions, liquid- or gel-like condensates can further transition into an aggregated or solid-like phase, characterized by stable interactions. The resulting aggregates have been linked to various pathological conditions, including neurodegenerative diseases such as Huntington’s disease, Alzheimer’s disease, amyotrophic lateral sclerosis, and frontotemporal dementia^11,12^.

One example of a group of IDPs includes the FG-Nucleoporins (FG-Nups) of the nuclear pore complex (NPC) that are characterized by an enrichment in phenylalanine-glycine (FG) repeats. FG-Nups have a folded domain to anchor them to the NPC and an IDR encoding the FG-repeats. Eleven different FG-Nups, each present in multiple copies, make up the ∼200 FG-Nups that project into the NPC’s channel in yeast cells, and similar numbers apply for the human NPC^13–15^. Together with nuclear transport receptors, the FG-Nups form the selective permeability barrier, which regulates all selective nucleocytoplasmic transport of macromolecules^16–19^. The FG-Nups anchored inside the NPC are highly dynamic^20^, but outside of the NPC context, FG-Nups or fragments of FG-Nups encoding only the ID region can spontaneously undergo phase separation into liquid-, gel-, or amyloid-like states^21–26^. Nup condensates can be functional and have been described as storage and assembly platforms in Drosophila oogenesis, in HeLa cells, and in *Saccharomyces cerevisiae*^27–32^. FG-Nups also appear in dysfunctional condensates and aggregates in disease contexts^33–39^. Their ID sequences are exemplary of an organization in “stickers and spacers,” where stickers represent the regions or residues that primarily drive attractive multivalent interactions, whereas spacers tune the solubility and regulate the phase behavior of IDPs^40,41^. Related to the differences in amino acid sequence (e.g. the type and density of FG repeats, position and presence of charged residues), some FG-Nups form condensates more readily than others^21–26,13,42^. For example, FG-Nups with GLFG repeats have a stronger tendency to self-associate into condensates than other FG-Nups^13,21,26,42^. The mixture of FG-Nups present in the NPC and in natural condensates has been proposed to be evolutionary tuned to stabilize each other, where the FG-Nup Nsp1 is a strong modulator of FG-Nup condensates, promoting a liquid-like state^29^.

We^43,44^ and others^45^ previously showed that the molecular chaperone DNAJB6 can directly interact with FG-Nups and delay their transition to more solid-like FG-Nup particles. We termed this phase-state surveillance. This new discovered function is relevant for NPC assembly^43,45^, and adds to its well-established role in interacting with and delaying the aggregation of several other IDPs (polyQ, FUS, FTD) *in vitro*, in cells, and in animal models^43,46–57^. DNAJB6 is a class B’ J-domain protein (JDP) and human cells express two alternatively spliced isoforms: the longer, nuclear isoform a (DNAJB6a), and the shorter isoform b (DNAJB6b), which is found both in the nucleus and the cytoplasm, and is the one referred to in this paper. In DNAJB6, the N-terminal J-domain is followed by a glycine (G) and phenylalanine (F) rich region (G/F region) and a serine (S) and threonine (T) rich region (S/T region) that is also rich in F and G residues. Previous studies showed that the S and T residues in the S/T region are crucial for DNAJB6’s ability to delay the aggregation of polyQ^46,53^, while the F residues in the same region are crucial for the anti-aggregation activity towards FG-Nups^43^. Notably, the G/F and S/T regions of DNAJB6 are predominantly disordered, except for a short regulatory helix (residues 95-104) known to modulate the interaction with HSP70^58,59^. The C-terminus of DNAJB6 consists of four β-strands and was shown to be crucial to inhibit secondary nucleation in amyloidogenesis^60^.

DNAJB6’s ortholog DNAJB8, which is typically expressed only in testis, and DNAJB2 isoform a (hereafter referred to as DNAJB2) also belong to the class B’ JDPs with a non-canonical architecture^61^ and have a similar IDR sequence composition (located between the regulatory helix and the C-terminal β-sheets), including the presence of predicted LARKS. Besides their similar IDR architecture, DNAJB6, DNAJB2, and DNAJB8 are all able to accumulate in nuclear envelope (NE) herniations^43,45^. These herniations are stalled pre-fusion NPC assembly intermediates that accumulate MLF2 and DNAJB6 within their lumen when fusion of both NE membranes is blocked^43,62,63^.

Previous data showed that DNAJB6 and FG-Nups form partially colocalized condensates^43,44^, but it was unclear whether and how the self-association properties of both DNAJB6 and FG-Nups are important for their interaction and DNAJB6’s phase-modulatory activity towards FG-Nups. To explore this question and to understand how the primary sequence of the DNAJB6-IDR encodes the self-assembly properties of the chaperone, we examined the activity of DNAJB6, DNAJB2 and DNAJB8 across a range of FG-Nup clients and conducted a targeted mutational and evolutionary analysis of the IDR of DNAJB6. We complemented our experimental in cell and *in vitro* analyses with multiscale molecular dynamics simulations aimed at probing the thermodynamic stability, intermolecular interaction landscape, and β-sheet-forming propensity of the wildtype and different scrambled variants of DNAJB6. Our analysis elucidates that the surveillance mechanism of DNAJB6 is encoded in this highly conserved, LARKS-rich disordered region which promotes the formation of stable gel-like chaperone assemblies.

## Results

### DNAJB6 forms gel-like assemblies in vitro and in cells

Previous data showed that purified DNAJB6 forms self-associated assemblies, which, depending on the conditions, range from ∼25 nm^47,64–66^ to µm-sized^43,44,57^. The latter larger assemblies form *in vitro* after 1 hour at 3 µM concentration and in buffers with or without 10% of the molecular crowder PEG. They appear as heterogeneous grape-like assemblies that were not observed to fuse or split and that partly (54%) remain after a 10-minute exposure to 5% 1,6-hexanediol (**Figure 1A**). Fluorescence Recovery After Photobleaching (FRAP) experiments in HEK293T^WT^ cells overexpressing mCherry-DNAJB6 concur with this (**Figures 1B,C)**. We find that the spherical assemblies that form upon overexpression of the GFP-tagged FG-rich region of Nup153 (GFP-Nup153FG) exhibit fast recovery kinetics (t½ ≈ 8.1 seconds) and a high mobile fraction (97%). In contrast, when co-localized with mCherry-DNAJB6, the recovery of GFP-Nup153FG was significantly delayed (t½ ≈ 9.8 seconds) and the mobile fraction was reduced (67%). Strikingly, the recovery of mCherry-DNAJB6 within these same assemblies was markedly slower (t½ ≈ 19.6 seconds) and remained incomplete within the acquisition time window. The DNAJB6 assemblies formed in cells thus slow down both internal molecular mobility of FG-Nups and exchange with the surrounding environment.

**Figure 1.**
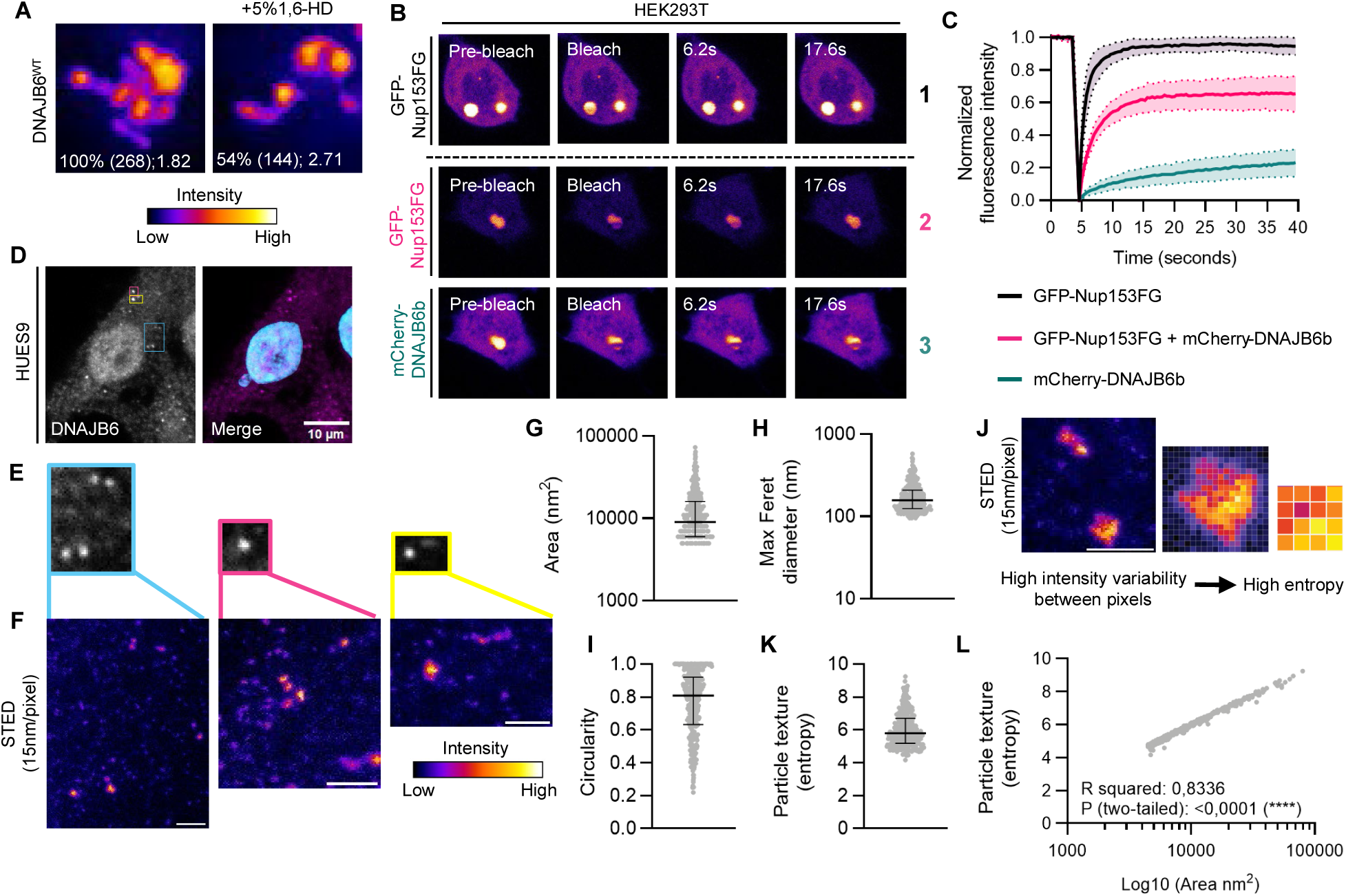
DNAJB6 forms gel-like assemblies *in vitro* and in cells. **(A)** Representative images of *in vitro* purified DNAJB6^WT^ particles formed in the presence of 10% PEG3350 (1h) and displayed according to fluorescence intensity. After 1 hour incubation, samples were treated with 5% 1,6-hexanediol for 10 minutes. Indicated are: % particles remaining (total nr particles); median particle size in µm (n=2) **(B)** Representative images of Fluorescence Recovery After Photobleaching (FRAP) experiments performed in HEK293T^WT^ co-transfected with GFP-Nup153FG (encoding Nup153_875-1475_) in the absence or presence (below dashed line) of mCherry-tagged DNAJB6b. **(C)** Normalized FRAP curves from the GFP channel of GFP-Nup153FG particles alone (condition 1, black) or co-localized with mCherry-DNAJB6b (condition 2, pink), and from the mCherry channel of GFP-Nup153FG particles co-localized with mCherry-DNAJB6b (condition 3, teal). Graphs show mean ± SD of three independent experiments with N=39 particles for condition 1 and N=35 particles for conditions 2 and 3. **(D)** Representative confocal image of a human embryonic stem cell (ESC) stained for endogenous DNAJB6 and Hoechst 33342. **(E)** Zoomed-in panels of three distinct cytoplasmic regions showing DNAJB6 foci. **(F)** STED images of the DNAJB6 foci shown in (D), displayed according to fluorescence intensity. Scale bar: 500 nm. **(G-I)** Area (G), maximum Ferret diameter (H), and circularity (I) of cytoplasmic DNAJB6 assemblies exemplified in (F). **(J)** Schematic of particle texture analysis. Left: representative STED image of two cytoplasmic DNAJB6 assemblies colored by fluorescence intensity. Scale bar: 500 nm. Center: zoomed-in assembly showing individual pixel intensities. Right: magnified 4×4-pixel region illustrating high local intensity variability. **(K,L)** Particle texture (entropy) (K) and correlation between particle texture (entropy) and area (L) for the 359 cytoplasmic DNAJB6 assemblies quantified in (G-I).

We assessed whether similar assemblies of DNAJB6 form endogenously in human embryonic stem cell (HUES9) line. Strikingly, regular confocal microscopy showed very frequent cytoplasmic DNAJB6 assemblies that appeared heterogeneous and amorphous **(Figures 1D,E)**. To better assess their features, we performed super resolution Stimulated Emission Depletion (STED) microscopy **(Figure 1F)**, which confirmed that DNAJB6 forms heterogeneous, irregularly shaped assemblies (**Figures 1G-I**). Additionally, texture analysis revealed a non-homogeneous internal signal distribution in these assemblies, with high entropy values that scale proportionally with particle area **(Figures 1J-L)**, suggesting cytoplasmic DNAJB6 might organize into heterogeneous, structurally complex assemblies. To our knowledge, this is the first demonstration that endogenous DNAJB6 organizes into cytoplasmic assemblies under physiological conditions.

Altogether, we conclude that DNAJB6 forms heterogeneous, irregularly shaped assemblies that have characteristics of being gel-like assemblies both *in vitro,* in cells when overexpressed, and, importantly, also endogenously in the cytoplasm of Hues9 cells.

### DNAJB6 can modulate a range of NupFG condensates

To begin to address the question if the activity of DNAJB6 in surveilling FG-Nups relates to its propensity to form gel-like assemblies, we first addressed how the surveillance of DNAJB6 depends on the condensation of the FG-Nups. We used a panel of purified nucleoporin fragments encoding the ID region of the FG-Nups and we refer to them as NupFG (see methods for precise regions). We systematically analyzed the size, circularity, and intensity of large numbers of Nup60FG, Nup100FG, Nup116FG, Nup145NFG and Nup153FG particles formed in the presence and absence of DNAJB6. We used two buffers, with and without the molecular crowder PEG (10%) and imaged at two timepoints, 1 hour and 24 hours (**Figure 2A**). DNAJB6 itself also forms particles under the conditions tested (**Figure S1A**).

**Figure 2.**
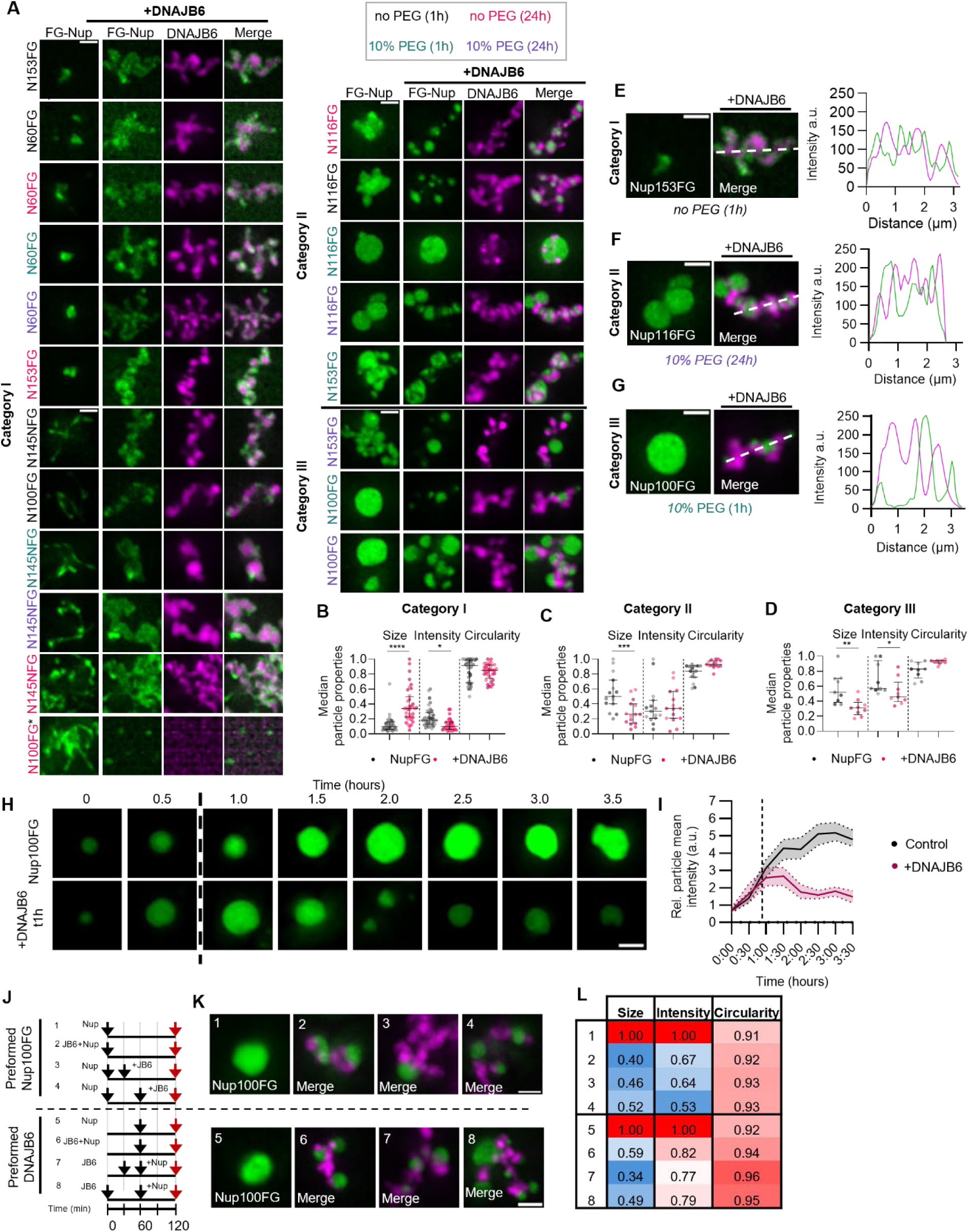
DNAJB6 can modulate a range of NupFG condensates. **(A)** Representative images showing Nup60FG-5MF, Nup100FG-5MF, Nup116FG-5MF,Nup145FG-5MF and Nup153FG-5MF particles, formed in the absence or presence of 10% PEG3350 for 1h or 24h, in the absence or presence of DNAJB6-A594 (molar ratio 1:1). The total set was sorted and subdivided into three categories, based on the particle properties of the control condition and the type of mixtures formed between the different FG-Nup fragments and DNAJB6. Representative images and categories were based on quantitative analysis of >300 particles per condition (n=4). For the condition indicated by * a second phenotype was observed; See Figure S1B. Scale bar, 1 µm. **(B-D)** Mean size, fluorescence intensity and circularity of particles belonging to category 1 (B), category 2 (C) and category 3 (D). See Figures S1I-N for quantifications; >300 particles per condition (n=4). **(E-G**) Exemplary particle belonging to category 1-3, respectively, alongside line scan profile. Quantified overlapping areas are found in (Figure S1D). Scale bar, 1 µm. **(H)** Representative images of Nup100FG-5MF particles formed in the presence of 10% PEG3350 and followed over time (in hours). Time of addition of DNAJB6 is indicated by the black arrow. Scale bar, 1 µm. **(I)** Mean fluorescence intensity of Nup100FG-5MF particles exemplified in (H), relative to the mean of the control at t0. The black-dotted lines indicate the time of addition of DNAJB6b. Graph shows the median ± 95% CI of all measurements from two replicates. 100 particles were analysed for each time point for each of the independent replicate experiments. **(J)** Schematic diagram of mixing conditions for experiments depicted in (K). Red arrow indicates time of imaging. **(K)** Representative images showing Nup100FG-5MF particles in the absence or presence of DNAJB6-A594 (molar ratio 1:1). Numbers refer to conditions in (J). Scale bar, 1 µm. **(L)** Table showing overview of median particle size, intensity and circularity for each of the conditions exemplified in (K). A colour-gradient was applied to each of the assessed particle properties, representing the magnitude of change relative to the control condition, with lower values indicated in blue and higher values indicated in red.

When comparing the properties of the NupFG particles formed in the absence and presence of DNAJB6, we discriminate three different behaviors. A first group of NupFGs (**Figure 2A**, category 1), which themselves only form few small round or irregularly shaped particles, bind to the larger DNAJB6 particles resulting in a change in the NupFGs particle properties: they become larger, less intense, and less circular and display a median overlap of 91% between the NupFG and DNAJB6 signals (**Figures 2B, E**; quantification of individual FG-Nups in **Figure S1C**). A second group of NupFGs (**Figure 2A**, category 2), which form larger particles in the absence of DNAJB6, also showed high overlap with DNAJB6 (median 90%), and these particles became smaller and more circular in the presence of DNAJB6 (**Figures 2C,F; Figure S1C**). A third group of NupFGs (**Figure 2A**, category 3) is rapidly condensating, forming large and intense particles on their own. In the presence of DNAJB6 these NupFG particles become smaller, less intense, and more circular (**Figures 2D,G; Figure S1C**), and only show small regions of overlap with DNAJB6 (median 45%). Rather, DNAJB6 appears to wrap around these condensates, forming small contact sites at the surface, with some partial incorporation into the NupFG condensates. Combined, DNAJB6 can engage with all NupFG condensates, albeit in different manners depending on the NupFGs particle properties, indicative of their phase state^44^.

Having observed that DNAJB6 can engage with a wide range of NupFG condensates with distinct particle properties, we aimed to test whether DNAJB6 could also engage with preformed NupFG particles. For this, we followed the dynamics of Nup100FG particles in time (**Figure 2H**). As expected, we see that with time the particles increase in intensity and size. The addition of DNAJB6 after one hour resulted in a decrease in particle intensity (**Figures 2H-I**), confirming that DNAJB6 can also act on pre-formed Nup100FG particles. We tested several schemes of preforming Nup100FG or DNAJB6 particles before mixing (**Figure 2J**) and assessed the properties of the Nup100FG particles (**Figure 2K-L**) and the appearance of a 0.5% SDS-insoluble Nup100FG fraction which can be trapped in filter trap assays (FTA) (**Figures S1E-H**)^43^. Both assays revealed that Nup100FG particles showed a similar reduction in intensity and size and increase in circularity (**Figure 2L**, **Figures S1I-N**), and a similar reduction of Nup100FG aggregation, regardless of the timing of their mixing (**Figures S1E-H**).

Our systematic assessment of the condensation behavior of different NupFG fragments in combination with DNAJB6 shows that DNAJB6 interacts with all NupFG particles under all conditions, suggesting that the activity of DNAJB6 is not limited to a specific NupFG phase state. The ability of DNAJB6 and Nup100FG to efficiently engage in heterotypic interactions even when either of these proteins is preassembled further highlights their capacity to outcompete homotypic interactions and engage in heterotypic interactions. Our data align well with recent studies reporting that DNAJB6 and FUS co-phase separate, forming a loose gel-like state which delays FUS fibrilization^57^.

### The phenylalanine residues in the S/T-rich region of DNAJB6 are essential for its ability to self-associate and modulate FG-Nup phase transitions

In our previous work, we showed that the S/T region of DNAJB6 is important for both its anti-amyloidogenic effects on polyQ^46,53^ as well as for its action to prevent liquid-to-solid transitions of FG-Nups^43^. The canonical type B JDP, DNAJB1, that lacks such a region **(Figures 3A,B)**, can disaggregate polyQ as a trimeric complex together with HSP70 and HSP110^67–69^, but it cannot independently act on polyQ fragments^47^. Here, we show that purified DNAJB1 is also not effective at delaying the aggregation of FG-Nups (**Figures 3C,D**). However, when inserting the S/T region from DNAJB6 C-terminally to the G/F-rich region of DNAJB1 (DNAJB1^DNAJB6-S/T^) (**Figures 3A,B**), it does gain anti-aggregation activity towards Nup100FG, albeit not to the same extent as wildtype DNAJB6 (DNAJB6^WT^) (**Figures 3C,D**). These results further highlight that the S/T region is critical for chaperoning FG-Nups.

**Figure 3.**
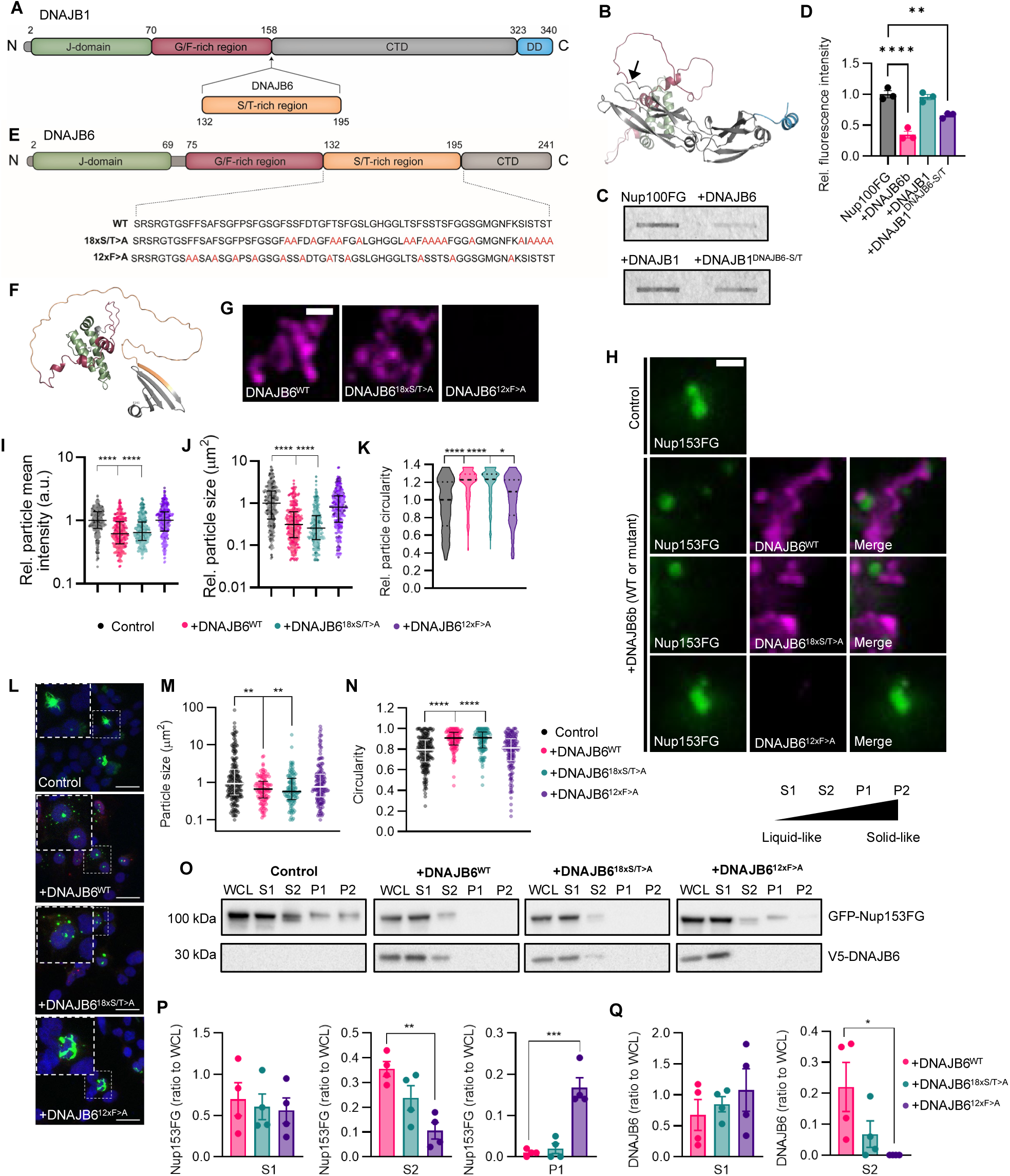
The phenylalanine residues in the S/T-rich region of DNAJB6 are essential for its phase modulatory activity. **(A)** Schematic overview of the domain structure of DNAJB1, indicating the N-terminal J domain (green), followed by the G/F,G-M domain (pink), the the C-terminal domain (CTD) (grey) and the DD domain (blue). The S-T domain of DNAJB6 (yellow) is inserted immediately after the G/F-rich domain. **(B)** Predicted structure of DNAJB1 by AlphaFold ^105^, with the arrow indicating the location where the S/T-rich domain of DNAJB6 is inserted. **(C)** Filter trap assay to assess aggregated fraction of Nup100FG-5MF in the absence or presence of either DNAJB6^WT^, DNAJB1^WT^ or DNAJB1^DNAJB6-S/T^. **(D)** Quantification of the band intensities of Nup100FG-5MF on filter trap. Represented band intensities are relative to the average intensity of the control. Mean ± SEM (n=3). **P<0.01, ***P<0.001. **(E)** Schematic overview of the domain structure of DNAJB6b, indicating the N-terminal J domain (green), followed by the G/F domain (pink), the S/T domain (yellow) and the C-terminal domain (CTD) (grey). The locations of the different mutations (18×S/T > A, 12×F > A) are indicated. **(F)** Predicted structure of DNAJB6 by Alphafold. **(G)** Representative images of wildtype and indicated DNAJB6 mutant (18x S/T > A, 12x F > A) particles, formed in the presence of 10% PEG3350 (1h). Scale bar, 1 µm. **(H)** Representative images showing Nup153FG-5MF particles in the absence or presence of either DNAJB6^WT^, DNAJB6^18xS/T>A^ or DNAJB6^12xF>A^ particles, formed in the presence of 10% PEG3350 (1h) (molar ratio 1:1). Scale bar, 1 µm. **(I-K)** Mean fluorescence intensity (I), size (J) and circularity (K) (relative to the median of the control) of Nup153FG-5MF particles exemplified in **(H)**. Graphs show median ± interquartile range of ≥300 particles per condition (n=3). **(L)** Representative images of HEK293T^DNAJB6-/-^ co-transfected with GFP-Nup153FG in the absence or presence of V5-tagged DNAJB6b constructs (DNAJB6^WT^, DNAJB6^18xS/T>A^, or DNAJB6^12xF>A^). Scale bar, 20 µm. **(M,N)** Size (M) and circularity (N) of Nup153FG particles exemplified in (L). Graphs show median ± interquartile range of combined particles from 3 independent replicates. **(O)** Representative western blot showing the different fractions from a protein fractionation assay performed in HEK293T^DNAJB6-/-^ cells co-transfected with GFP-Nup153FG in the presence or absence of V5-tagged DNAJB6b constructs (DNAJB6^WT^, DNAJB6^18xS/T>A^, or DNAJB6^12xF>A^). WCL: whole cell lysate. S1: 0.5% NP40 soluble. S2: 0.1% SDS soluble. P1: 2% SDS soluble. P2: 2% SDS insoluble. **(P)** GFP-Nup153FG band quantification in the different fractions and conditions expressing V5-tagged DNAJB6b shown in (O). Graphs show mean ± SEM (n=4). **(Q)** V5-DNAJB6 band quantification in the different fractions and conditions expressing V5-tagged DNAJB6b shown in (O). Graphs show mean ± SEM (n=4). *P<0.05, **P<0.01, ***P<0.001, ****P<0.0001.

The S/T region is enriched in S, T, and F residues that, in cellular experiments, were shown to be essential for the chaperoning capacity of DNAJB6^24,27^. To get insight in their loss of function features we purified mutants in which either 18 S and T residues were changed to alanine (DNAJB6^18xS/T>A^) or all 12 F residues were changed to alanine (DNAJB6^12xF>A^) (**Figures 3E,F**). We found that purified DNAJB6^18xS/T>A^ still self-associates, whereas DNAJB6^12xF>A^ completely lost this ability (**Figure 3G**). In line, DNAJB6^18xS/T>A^ still colocalizes with Nup153FG particles and reduces their aggregation propensity (**Figures 3H-K**), similar to DNAJB6^WT^. In contrast, purified DNAJB6^12xF>A^ was unable to influence the size, intensity, or circularity of Nup153FG particles (**Figures 3H-K**), consistent with previous cellular data^43^. Consistent results were obtained for the interaction of both mutants with Nup100FG (**Figures S2A-D**).

To corroborate these findings in cells, V5-tagged versions of both mutants were co-overexpressed with GFP-Nup153FG in DNAJB6 knockout cells (HEK293T^DNAJB6–/–^). The size and circularity of the GFP-Nup153FG particles, as well as their detergent solubility were analyzed via microscopy and protein fractionation assays, respectively. Whereas overexpression of DNAJB6^18xS/T>A^ suppressed the formation of large and irregularly-shaped (possibly amyloidogenic) GFP-Nup153FG particles, overexpression of DNAJB6^12xF>A^ was unable to do so (**Figures 3L-N**). Similarly, DNAJB6^18xS/T>A^, but not DNAJB6^12xF>A^, suppressed the accumulation of GFP-Nup153FG in the SDS-insoluble fraction (P1) as efficiently as DNAJB6^WT^ (**Figures 3O,P**). In line with our *in vitro* data, in cells DNAJB6^12xF>A^ also remains fully soluble, while a fraction of DNAJB6^WT^ is found in particles that are resistant to 0.5% NP40 (S2) (**Figures 3O,Q**). So, also in cells the F residues in the S/T region of DNAJB6 are essential for its ability to self-associate and modulate FG-Nup condensation and aggregation.

To assess whether these two mutants impair DNAJB6’s function related to NPC assembly, we assessed their ability to accumulate in NE herniations. We quantified the co-localization of MLF2 (a herniation marker^63^) and DNAJB6 foci in HEK293T^WT^ cells overexpressing the different V5-tagged DNAJB6 constructs. Immunofluorescence experiments revealed that DNAJB6^18xS/T>A^, but not DNAJB6^12xF>A^, still accumulates into NE herniations (**Figures S2E,F**). Thus, DNAJB6’s ability to phase separate and delay the aggregation of FG-Nups is connected to its accumulation in NE herniations and, potentially, its function in NPC assembly.

Combined, our data suggests that DNAJB6’s ability to interact with and delay the phase transitions of its various FG-Nup substrates depends on the phenylalanine residues in its S/T region and its self-association ability.

### Specificity of DNAJB2, DNAJB6, and DNAJB8 for FG-Nups

We next assessed whether the ability to interact with and modulate the phase state of a range of FG-Nups was particular to DNAJB6 or whether similar JDPs would share this ability. Bioinformatics analysis showed that the IDRs of the class B’ JDPs^61^ (DNAJB2, DNAJB6, DNAJB7, and DNAJB8) located between the regulatory helix and the C-terminal β-sheets (IDR1), are also enriched in F residues, similar to some FG-Nups (**Figures 4A,B**). Using the NARDINI algorithm^70^ to assess the binary amino acid distribution and content in a given sequence, we find that there are minor differences in their polar and glycine residue distribution, but they all exhibit a random arrangement of aromatic residues (**Figure S2G**). Clustering analysis showed that the IDR1 from DNAJB2, DNAJB6, and DNAJB8, but not DNAJB7, closely clustered together based on their amino acid distribution and composition (**Figure S2G**), which prompted us to test whether DNAJB2 and DNAJB8 could also display phase modulatory activities towards different FG-Nups.

**Figure 4.**
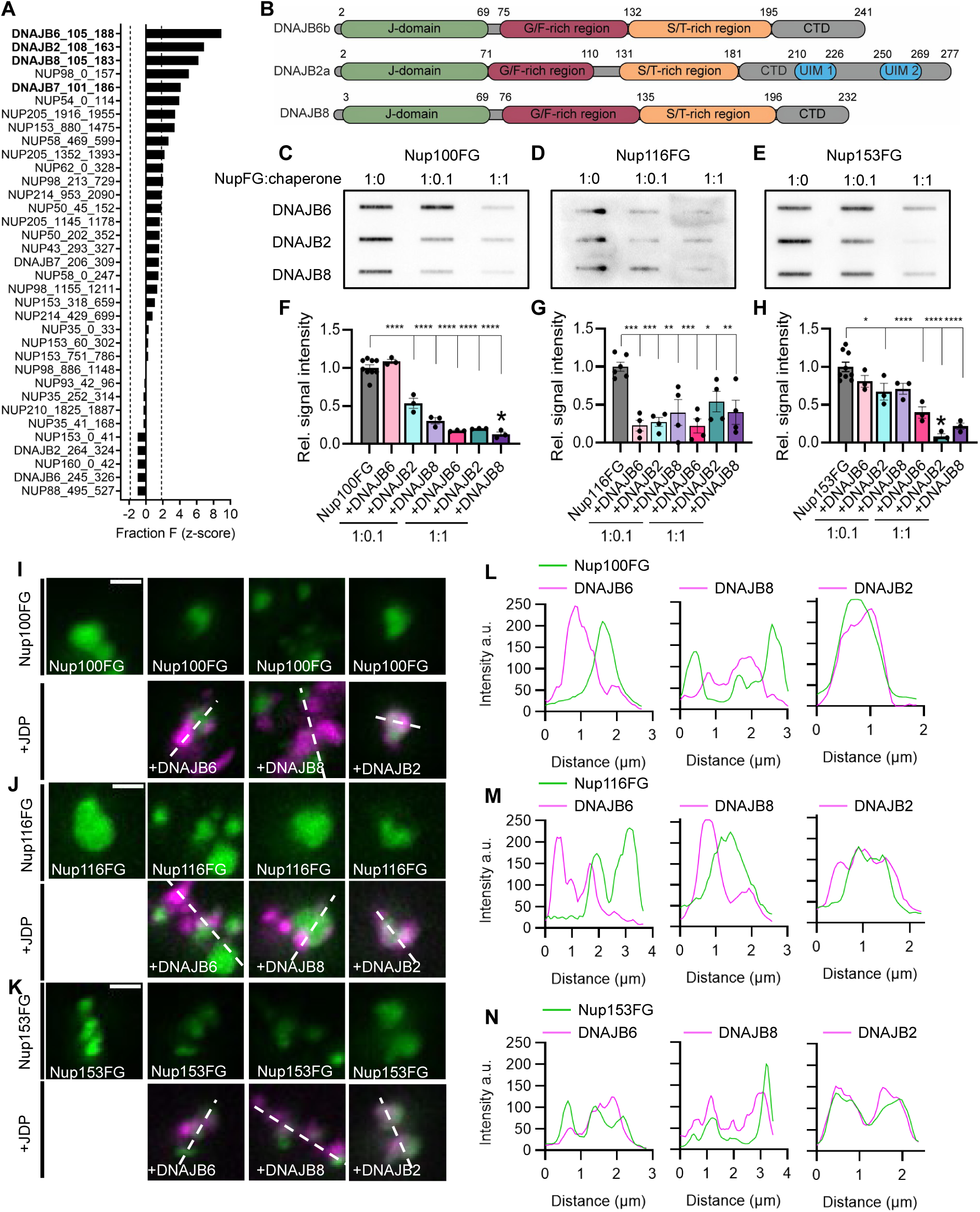
Specificity of DNAJB2, DNAJB6, and DNAJB8 for FG-Nups. **(A)** Z-scores for phenylalanine (F) fractions in the IDRs of class B’ JDPs and human FG-Nups. Dotted lines indicate the -1.96 and +1.96 boundaries for the random distribution. The IDRs from class B’ JDPs located between the regulatory helix V and the C-terminal β-sheets are highlighted in bold. **(B)** Schematic overview of the domain structure of DNAJB6, DNAJB2 and DNAJB8, indicating the N-terminal J domain (green), followed by the G/F domain (pink), the S/T domain (yellow) and the C-terminal domain (CTD) (grey). **(C-E)** Filter trap assay to assess aggregated fraction of Nup100FG (C), Nup116FG (D) and Nup153FG (E) in the absence or presence of either DNAJB6, DNAJB2 or DNAJB8 at a molar ratio of either 1:10 or 1:1 (Nup100FG (3µM), Nup153FG (6µM) (1h), Nup116FG (6µM) (3h)). **(F-H)** Quantification of the band intensities of Nup100FG (F), Nup116FG (G) and Nup153FG (H) on filter trap. Represented band intensities are relative to the average intensity of the control. Mean ± SEM (n=3). *P<0.05, ****P<0.0001. **(I-K)** Representative images showing Nup100FG-5MF (I), Nup116FG-MF (J) and Nup153FG-MF particles (K), in the absence or presence of either DNAJB6b, DNAJB8 or DNAJB2a particles, formed in the presence of 10% PEG3350 (1h) (molar ratio 1:1). Scale bar, 1 µm. **(L-N)** Line scan profiles showing interaction profiles for representative images of Nup100FG (L), Nup116FG (M), and Nup153FG (N) in the presence of DNAJB6, DNAJB8 and DNAJB2, respectively.

Like DNAJB6, both delayed the aggregation of Nup100FG, Nup116FG and Nup153FG (**Figures 4C-H**) with some specificity: while DNAJB8 was found to be most effective against aggregation of Nup100FG at both tested ratios (NupFG:chaperone 1:0.1 and 1:1) (**Figures 4C,F**), DNAJB2 was found to be most effective against aggregation of Nup153FG (**Figures 4E,H**). Microscopy assessment showed that whilst DNAJB6 and DNAJB8 wrap around NupFG condensates forming small contact sites at the surface, or partially incorporate into Nup116FG and Nup153FG condensates, DNAJB2 fully partitions into all FG-Nup condensates (**Figures 4I-N**). Combined, these findings indicate that DNAJB6, DNAJB8, and DNAJB2 all possess the capacity to self-associate and to engage with FG-Nups to modulate their phase state, but with some degree of substrate specificity.

### Scrambling the primary amino acid sequence of DNAJB6-IDR affects the solubility and self-association behavior of DNAJB6

In addition to the specific amino acid composition, the particular distribution of amino acids within IDPs has been demonstrated to influence both condensation properties and function^71^. In line with this, we aimed to determine the sequence features that may dictate the self-association behavior and phase modulatory activity of DNAJB6. We notice that the IDR from DNAJB6 (residues 105-108; DNAJB6-IDR) displays six binary combinations from its polar, aromatic, and glycine residues, which do not organize in any specific pattern, but are rather randomly distributed (**Figure S2G**). A binary amino acid distribution is random if the z-score falls between -1.96 and +1.96, segregated if > +1.96, or uniformly distributed if < -1.96. Of note, the DNAJB6-IDR also comprises a few charged residues that concentrate in the beginning of the sequence that are just below the 10% threshold from the NARDINI algorithm, and their distributions are, therefore, not computed.

To probe whether maintaining such random distribution is sufficient to maintain the self-association behavior and activity of DNAJB6, we used the NARDINI algorithm to generate three scrambled sequences (SCR1, SCR2, and SCR3) with a z-score matrix that is most similar to the query sequence while less than 15% of the amino acids are kept in their original position (**Figures 5A,B**). Two additional mutants (rL1 and rL2) were generated from the disordered S/T region only (residues 136-188) without controlling for their random distribution (**Figure 5A**). In DNAJB6^rL1^ the residues were completely scrambled using a random sequence generator, which unintentionally introduced the clustering of polar-polar residues, while the other residues were kept randomly distributed (**Figures 5A,B**). In DNAJB6^rL2^ we aimed to test the importance of the pairing of F’s and G’s, a feature that is characteristic for FG-Nups, so we deliberately split them apart (**Figures 5A,B**). As expected, DNAJB6^rL2^ shows a significant segregation of aromatic and glycine residues. However, this splitting also influenced the other binary combinations involving aromatic and glycine residues, as evidenced by the segregation of polar-aromatic and aromatic-aromatic residues, and the uniform distribution of polar and glycine residues (**Figure 5B**).

**Figure 5.**
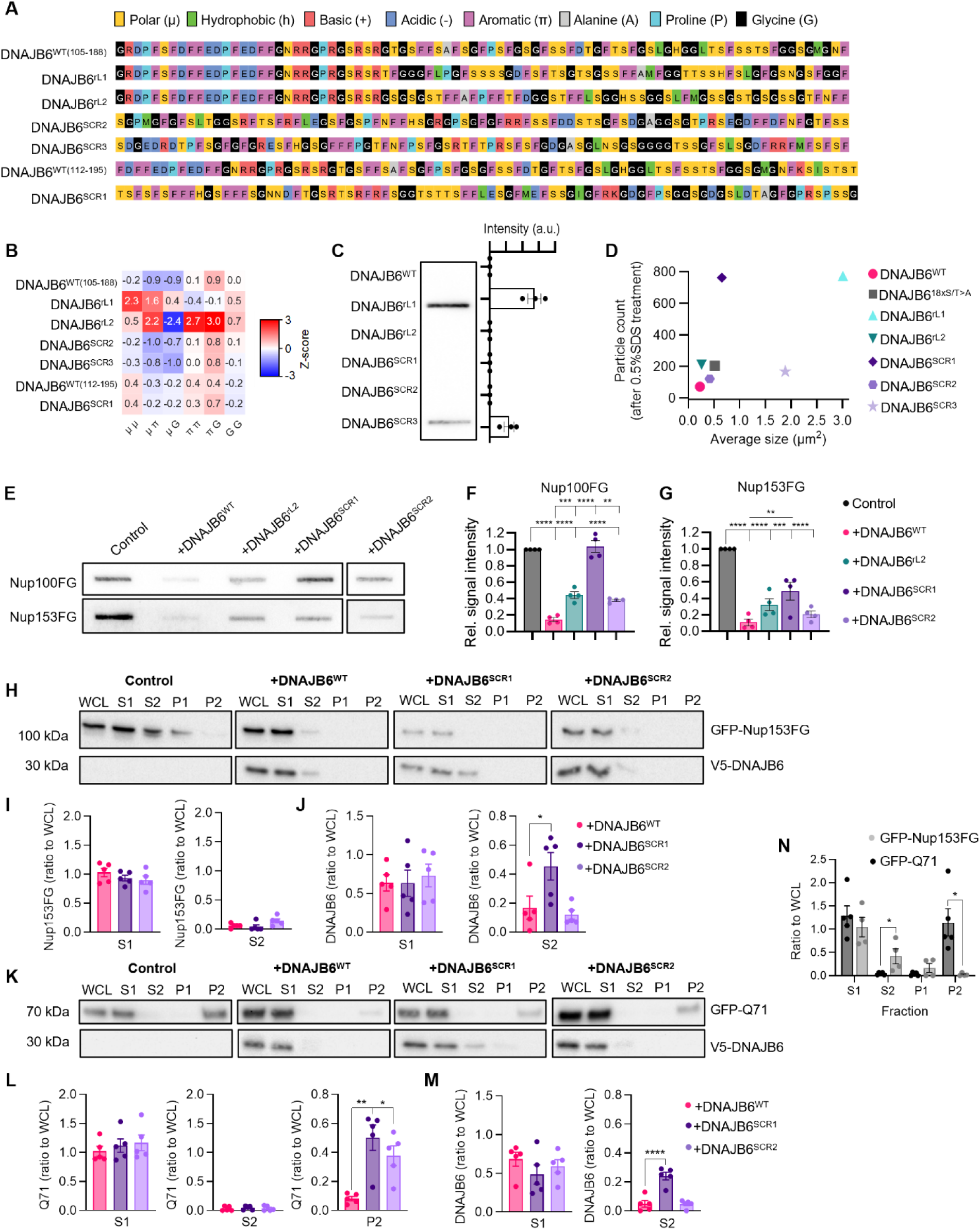
Scrambling the primary amino acid sequence of DNAJB6-IDR affects the solubility and activity of DNAJB6. **(A)** Primary amino acid sequences of DNAJB6^WT^ (residues 105-188 or 112-195) and mutants DNAJB6^rL1^, DNAJB6^rL2^, DNAJB6^SCR2^, DNAJB6^SCR3^ and DNAJB6^SCR1^ coloured by amino acid group. **(B)** Heatmap of z-score matrices from NARDINI algorithm showing binary amino acid distributions of the primary amino acid sequences shown in (A). Significantly positive (red; z-score > +1.96) values reflect that the pairs of types of amino acids have a non-random segregation from one another and from other residues, and significantly negative (blue; z-score < -1.96) values features non-random uniform dispersion of residues. Columns with null z-scores are omitted. **(C)** Filter-trap assay and quantification of band intensities to assess aggregated fraction of DNAJB6^WT,^ DNAJB6^rL1^, DNAJB6^rL2^, DNAJB6^SCR1^, DNAJB6^SCR2^, and DNAJB6^SCR3^ (n=3). **(D)** Quantification of particles formed by DNAJB6^WT^ and mutants (DNAJB6^18xS/T>A^, DNAJB6^rL1^, DNAJB6^rL2^, DNAJB6^SCR1-3^); representative images in **Appendix S3A**. (n=2). **(E)** Filter trap assay to assess aggregated fraction of Nup100FG-5MF and Nup153FG in the absence or presence of either either DNAJB6^WT^, DNAJB6^rL2^, DNAJB6^SCR1^ or DNAJB6^SCR2^. **(F,G)** Quantification of the band intensities of Nup100FG-5MF (F) and Nup153FG-5MF (G) on filter trap. Represented band intensities are relative to the average intensity of the control. Mean ± SEM (n=4). **(H,K)** Representative western blots showing the different fractions from a protein fractionation assay performed in HEK293T^DNAJB6-/-^ cells co-transfected with GFP-Nup153FG (H) or GFP-Q71 (K) and V5-tagged DNAJB6b constructs (DNAJB6^WT^, DNAJB6^SCR1^ or DNAJB6^SCR2^). WCL: whole cell lysate. S1: 0.5% NP40 soluble. S2: 0.1% SDS soluble. P1: 2% SDS soluble. P2: 2% SDS insoluble. **(I,L)** GFP-Nup153FG (I) or GFP-Q71 (L) band quantification in the different fractions and conditions expressing V5-tagged DNAJB6 shown in (H,K). **(J,M)** V5-DNAJB6 band quantification in the different fractions and conditions expressing V5-tagged DNAJB6 shown in (H,K). Graphs show mean ± SEM (n=4/5). **(N)** GFP-Nup153FG and GFP-Q71 band quantification in the different fractions of the control condition shown in (H,K). Graphs show mean ± SEM (n=4). *P<0.05, **P<0.01, ***P<0.001.

The five mutants were purified, and their particles were compared to those formed by DNAJB6^WT^. DNAJB6^SCR2^ and DNAJB6^rL2^ formed particles with similar size and sensitivity to treatment with 5% 1,6-hexanediol, and 0.5% SDS as the wildtype protein. The particles formed by DNAJB6^SCR3^ and, in particular, DNAJB6^rL1^ were however larger and partly resistant to 0.5% SDS (**Figures 5C,D, Figure S3A**). Imaging also showed some SDS-resistant DNAJB6^SCR1^ particles but those were small (**Figure 5D, Figure S3A**) and likely therefore not detected in the FTA (**Figure 5C**). We conclude that DNAJB6^rL1^ and DNAJB6^SCR3^ are particularly unstable (i.e. forming a 0.5% SDS-insoluble fraction) under the conditions tested while DNAJB6^SCR1^ is somewhat unstable. The observation that several of the mutants were less stable than the wildtype shows that maintaining a random amino acid distribution within the IDR of DNAJB6 does not suffice to maintain stable gel-like assemblies.

### Scrambling the primary amino acid sequence of DNAJB6-IDR more strongly impairs anti-aggregation of polyQ than FG-Nups

Next, we compared the functionality of the more stable DNAJB6^SCR2^, DNAJB6^rL2^ and partially unstable DNAJB6^SCR1^ mutant with DNAJB6^WT^. The DNAJB6 mutants have similar effects on the sizes and shapes of Nup100FG and Nup153FG particles compared to DNAJB6^WT^ (**Figures S3B-I**). Yet, biochemical assays showed a partial loss of function for DNAJB6^SCR2^ and DNAJB6^rL2^, and a more pronounced loss for DNAJB6^SCR1^, which was unable to reduce Nup100FG aggregation (**Figures 5E-G**). The mutants DNAJB6^SCR1^ and DNAJB6^SCR2^ were also assessed using the protein fractionation assay in HEK293T^DNAJB6-/-^ cell lysates. Cells overexpressing GFP-Nup153FG only showed accumulation of the NupFG in the P1 and P2 fractions, indicative of its aggregation (**Figure 5H**). In line with our previously published work^43^, the co-overexpression of V5-DNAJB6^WT^ largely prevented this (**Figures 5H,I**). Expression of either DNAJB6^SCR1^ or DNAJB6^SCR2^ delayed the aggregation of GFP-Nup153FG to the same extent, indicating that the partial loss of function observed *in vitro* was not detected in cells under the conditions tested (**Figures 5H,I**). Yet, we did observe that relatively more DNAJB6^SCR1^, but not DNAJB6^SCR2^, was found in the NP40-insoluble S2 fraction, potentially reflecting its altered self-association behavior observed *in vitro* (**Figures 5H,J**). So, DNAJB6^SCR1^, DNAJB6^SCR2^, and DNAJB6^rL2^ have altered self-association behavior with, as revealed *in vitro*, partially impaired functionality.

In the search for a substrate that might better reveal functional differences between DNAJB6^WT^ and the scrambled mutants in cells, we turned to testing their effect on the solubility of GFP-tagged Huntington exon 1 with a 71-polyQ stretch (GFP-Q71). This choice was instigated by our finding that DNAJB6^18xS/T>A^ showed unimpaired FG-Nup anti-aggregation activity (**Figure 3**), while it has reduced polyQ anti-aggregation activity^46^. In cells, the rate of liquid-to-solid phase transitions was faster for polyQ proteins than for FG-Nups, with polyQ appearing in the 2% SDS insoluble P2 fraction of control conditions (compare **Figures 5H,K**, quantified in **Figure 5N**). Interestingly, both the partially unstable DNAJB6^SCR1^ and the stable DNAJB6^SCR2^ showed impaired polyQ anti-aggregation activity compared to DNAJB6^WT^, as indicated by the increased aggregate levels in the P2 fraction (**Figures 5K,L**). Like in the experiments with Nup153FG (**Figure 5J**), relatively more DNAJB6^SCR1^ partitions to the S2 fraction (**Figure 5M**), underscoring that its altered stability is not related to the co-overexpressed substrate. Most importantly, these findings suggest that even for a mutant like DNAJB6^SCR2^, which forms similar assemblies as the wildtype DNAJB6, its phase state modulatory activity is partially impaired.

Together, our data points to a unique arrangement of amino acids in the DNAJB6-IDR which is essential for the stability of the chaperone and its anti-aggregation activity.

### The primary amino acid sequence of DNAJB6-IDR is highly conserved among mammals

Prompted by the idea that both the composition and arrangement of amino acids in the DNAJB6-IDR affect its stability and/or function, we decided to investigate the conservation of the primary amino acid sequence of this region in a curated dataset of 57 mammal species. For comparison, we included two closely related B’ class JDPs: DNAJB2 and DNAJB8, as well as an unrelated, but well-studied IDP: P53 (**Figure 6A**). The details on the selection of each protein’s IDR and folded domain are described in the methods section, and the DNAJB6-IDR is here referred to as IDR1. Multiple sequence alignments (MSAs) of the whole protein, folded domain, and IDRs were performed, and the residue conservation scores (maximum = 4.3, based on 20 amino acids) were calculated based on Shannon’s entropy, which quantifies variability at each alignment position. As expected, the folded domains showed the highest conservation scores, which were averaged to 4.0 and used as a visual reference score for highly conserved residues (**Figures 6B,C**). Interestingly, the IDR1 of DNAJB2, DNAJB6, and DNAJB8 showed significantly higher conservation compared to that of P53, and both DNAJB2 and DNAJB6 were notably more conserved in their IDR1 regions than DNAJB8 (**Figure S4C**). Given that DNAJB2 and DNAJB6 also contain a C-terminal IDR (IDR2), we included this region in our analysis for further comparison. Strikingly, the conservation scores for IDR2 were considerably lower than those for IDR1, with DNAJB6’s IDR2 being significantly less conserved than that of DNAJB2 (**Figure 6B** and **Figure S4D**). Aromatic and glycine residues were among the residues with the highest conservation scores in IDR1 of DNAJB2, DNAJB6, and DNAJB8 (**Figure 6D**). Consistent with IDRs generally showing conserved amino acid composition rather than position^72–76^, we find that this is also the case for the IDR1 of P53, DNAJB2, DNAJB6, and DNAJB8 (**Figures S4E-H**).These findings indicate that the IDR1 regions of DNAJB2, DNAJB6, and DNAJB8 are subject to stronger evolutionary pressures compared to a typical IDR, such as the one found in P53. For DNAJB6, this result corroborates the idea that maintaining the specific amino acid arrangement of IDR1 is crucial for achieving optimal stability and function.

**Figure 6.**
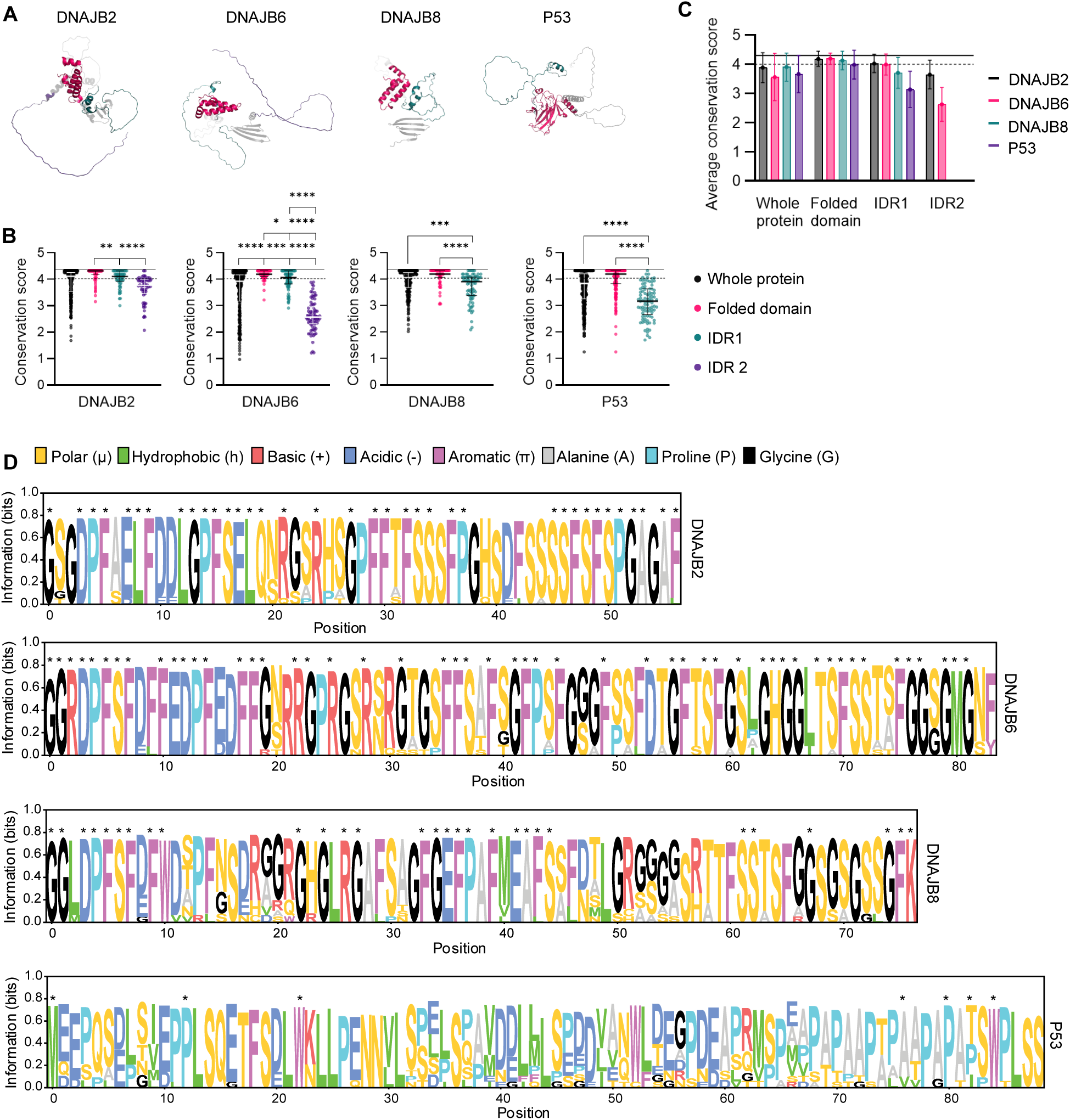
Evolutionary analysis of DNAJB6-IDR shows high primary amino acid sequence conservation. **(A)** Alphafold structures of DNAJB2, DNAJB6, DNAJB8, and P53 coloured by region (legend on panel C). Graph shows median with interquartile range. **(B)** Conservation scores from each region of DNAJB2, DNAJB6, DNAJB8, and P53. *P<0.05, **P<0.01, ***P<0.001, ****P<0.0001. **(C)** Conservation scores per region, per protein, from data shown in (B). Graph shows mean with 95% CI. Solid line at 4.3 indicates the maximum score based on Shannon’s entropy. Dashed line at 4.0 indicates average conservation score of all folded domains. **(D)** Sequence logo of IDR1 from DNAJB2, DNAJB6, DNAJB8 and P53 coloured by amino acid group. Asterisks (*) indicate residues with a conservation score > 4.

### Molecular simulations reveal LARKS-mediated tunning of the stability and material properties of DNAJB6 assemblies

To further investigate how sequence scrambling alters the physicochemical behaviour of DNAJB6 assemblies, we performed multiscale molecular simulations on DNAJB6^WT^ and its three scrambled variants (SCR1–3). We used a coarse-grained model (Mpipi-Recharged) designed to capture the condensate phase behaviour^77–79^. We first characterised the assembly formation of the four variants by means of direct coexistence simulations^78^, obtaining the temperature-concentration phase diagram (**Figure 7A**). According to the Mpipi-Recharged, SCR3 exhibited the highest critical solution temperature, indicative of enhanced assembly formation, whereas SCR1 showed the lowest critical temperature, consistent with a less cohesive phase. These differences are notable given the relatively modest sequence perturbations introduced by scrambling and agree with our biochemical observations showing SCR3 forms SDS insoluble fractions (**Figure 5C**). These results support the idea that the precise rearrangement of residues within the DNAJB6-IDR modulates assembly thermodynamics.

**Figure 7.**
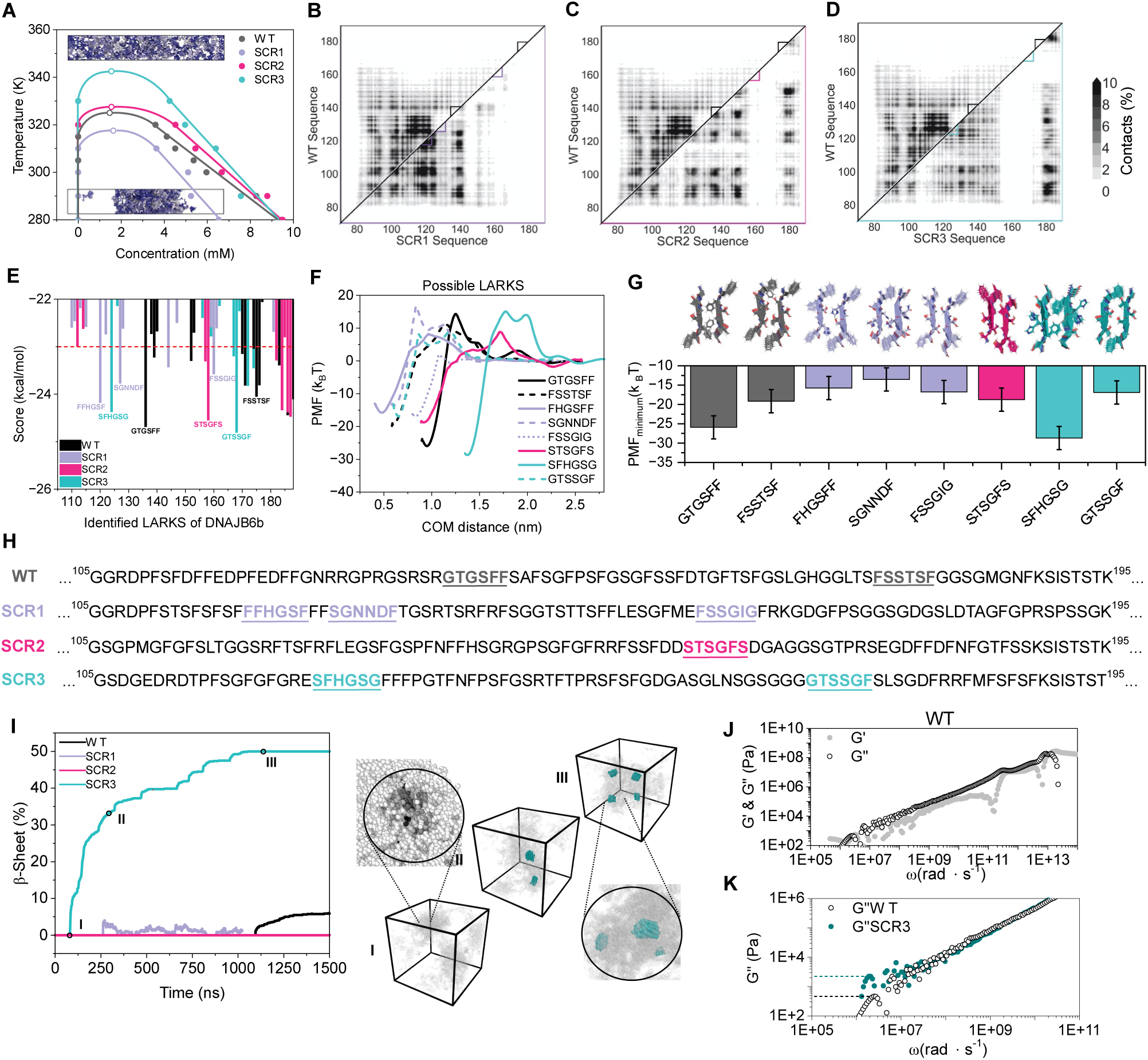
Coarse-grained simulations reveal that sequence scrambling alters intermolecular interactions, cross-β-sheet formation, and viscoelastic properties of DNAJB6b condensates. **(A)** Phase diagrams of WT DNAJB6b and its scrambled variants (SCR1, SCR2, SCR3) obtained via MD simulations. Representative simulation snapshots illustrate the two regimes: a homogeneous, single-phase system above the binodal (top), and a phase-separated configuration showing coexistence between a protein-dense condensate and a dilute phase (bottom). **(B-D)** Intermolecular residue–residue contact frequency maps (%) for the IDRs contained in the WT (top) and SCR1 (bottom) in (B), SCR2 (bottom) in (C) and SCR3 (bottom) in (D). All maps share a common colour scale with a maximum of 10% interaction frequency, where darker shading indicates higher contact probability. Triangular boxes highlight LARKS-containing regions. Amino acids are coloured according to their intermolecular contact frequency with residues belonging to other protein replicas. **(E)** ZipperDB energetic scores for LARKS segments identified along the DNAJB6b IDR sequence for each variant. The red dashed line denotes the energy threshold used to select candidate LARKS and evaluate them through all-atom simulations. See Figures S4L-P for individual scores. **(F)** Potential of mean force (PMF) as a function of center-of-mass (COM) distance for all selected LARKS across the different variants. **(G)** Minimum PMF values for each variant alongside representative structural snapshots of the two interacting fibril strands. **(H)** IDR sequences of each variant with the selected LARKS regions highlighted. **(I)** Cross-β-sheet formation kinetics expressed as the fraction of β-sheet strands (%) as a function of simulation time for each variant. Snapshots I, II, and III correspond to representative configurations of the SCR3 system at early, intermediate, and late stages of fibril assembly. **(J)** Viscoelastic moduli G′ and G″ as a function of frequency for the WT variant derived from the stress tensor. **(K)** Comparison of G″ in the low frequency regime between the WT and SCR3 sequences.

To further understand the molecular basis underlying the altered phase behaviour of the scrambled mutants, we next analysed the intermolecular residue–residue contact frequencies within the condensed phase for each sequence (**Figures 7B-D**, full contact map **Figures S4I-K**). In all DNAJB6 variants, intermolecular interactions were concentrated almost exclusively within the IDR, whereas the structured N- and C-terminal regions barely contributed to the assembly network connectivity. Importantly, the contact landscapes differed substantially between variants. Notably, in SCR3 and to a lesser extent in SCR2, strong intermolecular contacts contributing to phase-separation emerge between residues 178 and 185. Moreover, in SCR1 and SCR3 the strongest contact frequencies arise onto the aromatic-rich segments flanking LARKS motifs predicted by ZipperDB^80^ (depicted by small triangles across the diagonal for each variant and shown in **Figure 7E,H**, **Figures S4L,P**), suggesting that cooperative interactions between phenylalanine-rich regions and LARKS-mediated β-sheet interactions may stabilize intermolecular associations within the assembly. This observation aligns well with our experimental data showing that phenylalanine residues are essential for DNAJB6 self-association and phase-modulatory activity toward FG-Nups.

We next evaluated the energetic contribution of these LARKS motifs to form intermolecular cross-β-sheet interactions. To validate the binding energy of the predicted LARKS, we carried out atomistic potential of mean force (PMF) calculations^81–83^ which demonstrated that the strength of the intermolecular cross-β-sheet stacking differed substantially between variants (**Figures 7F,G**). In agreement with the weaker assembly stability observed for the SCR1 variant, its LARKS motifs displayed shallower free-energy minima, indicative of weaker and more reversible intermolecular interactions upon cross-β-sheet structural transitions. By contrast, SCR3 displayed interaction energies comparable to or stronger than WT, supporting more persistent cross-β-sheet assemblies. Since these calculations assess isolated LARKS motifs independently of the full sequence context, the enhanced aggregation propensity of SCR3 likely emerges from cooperative effects at the full-protein level, including altered frequencies and accessibility of interaction-prone regions.

To directly probe the consequences of these interaction networks on phase behaviour in the DNAJB6 assemblies, we monitored cross-β-sheet accumulation during coarse-grained simulations of assemblies formed by the different variants (**Figure 7I**). To that purpose, we used our previously developed out-of-equilibrium dynamic algorithm^84,85^ which mimics inter-protein β-sheet formation in a local-dependent and time-dependent manner for coarse-grained simulations. Whereas WT, SCR1, and SCR2 showed minimal or transient β-sheet formation within the assemblies, SCR3 rapidly accumulated stable β-sheet-rich assemblies over time. In SCR3, the β-sheet–forming LARKS corresponds to the SFHGSG motif, supporting our previous observations that also its surrounding chemical environment determines its ability to stabilize intermolecular contacts (**Figure 7D**). Interestingly, while the FHGSFF motif in SCR1 establishes a higher number of intermolecular LARKS–LARKS contacts than the corresponding motif in SCR3 (**Figures 7B,D**) its lower binding energy results in repeated formation and dissolution events, consistent with the observed transient cross-β-sheet formation. This behaviour is consistent with the presence of additional but energetically weaker LARKS motifs that increase nucleation frequency while failing to stabilize persistent fibrillar assemblies. Such transient interactions may help explain why SCR1 forms many small SDS-resistant particles experimentally (**Figure 5D, Figure S3A**), despite displaying only modest alterations in bulk assembly properties. A clear example reinforcing the importance of the local chemical and interaction environment within the sequence is SCR2, which, despite containing a LARKS motif with favorable intrinsic binding energy, fails to form fibrillar structures because the corresponding regions do not nucleate stable cross-β-sheet assemblies (see **Figure S4Q** for the LARKS contacts in the different variants and **Figure 7C**).

Finally, we assessed the material properties of the assemblies through rheological analysis^86^ (**Figure 7J,K**). While SCR1 and SCR2 behaved similarly to the WT sequence **(Figures S4R,S**), SCR3 exhibited a pronounced increase in the loss modulus at low frequencies (G″), consistent with enhanced viscoelasticity and a more viscous condensed phase. Together, these simulations provide a mechanistic framework that connects the experimentally observed changes in DNAJB6 stability and anti-aggregation activity to altered intermolecular interaction networks within the IDR. More broadly, these simulations support the notion that the highly conserved sequence organization of the DNAJB6-IDR is evolutionarily tuned to balance assembly stability, material properties, cross-β-sheet accumulation, and client surveillance. In this framework, aromatic interactions and LARKS-mediated β-sheet formation cooperate to stabilize DNAJB6 assemblies while limiting irreversible aberrant self-assembly.

## Discussion

DNAJB6 displays remarkable anti-aggregation activity towards several disease-related IDPs^46–49,51,53,54,56^, and recently has also been found to surveil the phase transitions of the intrinsically disordered FG-Nups^43–45^. When purified or when overexpressed in cells, DNAJB6 forms larger gel-like assemblies and here we show that also endogenous DNAJB6 forms assemblies in the cytosol of human embryonic stem cells.

We uncovered that the surveillance mechanism of DNAJB6 towards multiple FG-Nups is encoded in a surprisingly highly conserved, disordered region which promotes the formation of stable gel-like chaperone assemblies. Such assemblies provide an environment that enable low-affinity, dynamic heterotypic interactions, most likely between the phenylalanines in the DNAJB6-IDR and those in the FG-Nups. We argue these heterotypic interactions outcompete the homotypic FG-FG interactions of substrates, thereby delaying their aggregation. The evolutionary conservation of the DNAJB6-IDR, and our mutant analyses suggest that the sequence space for encoding stable gel-like assemblies is limited and finely tuned to balance a high local avidity while avoiding self-aggregation of the chaperone. Of note, the precise size and process underling the formation of such chaperone assemblies in the physiological cellular context remains to be resolved. Our work contributes to the view that molecular chaperones, beyond their classical role in protein (re)folding, are regulators of condensate dynamics.

### Gel-like DNAJB assemblies offer a protective environment against FG-Nup aggregation

Previous work showed that the disordered S/T region of DNAJB6 is critical for its activity towards FG-Nups and polyQ^43,46^. Here we substantiate this by showing that engineering the S/T region into DNAJB1 creates a hybrid chaperone (DNAJB1^DNAJB6-S/T^) that gains activity towards Nup100FG. Moreover, we showed that the F residues in this region are essential for its self-association into a gel-like state. This is consistent with recent findings from the Linse Group^87^ showing that this region, and not the β-sheet in the CTD^88^, is the primary driver of the self-assembly of full-length DNAJB6.

Our systematic analysis indicates that DNAJB6 interacts with and displays phase modulatory activity towards many FG-Nups with different condensation properties, both those that rapidly form gel-like condensates and those that are more soluble. DNAJB6 can even modify pre-assembled Nup100FG condensates making them smaller. The mechanism might involve disrupting the FG-Nup:FG-Nup intermolecular interactions, and/or acting as a physical barrier to prevent the coalescence and subsequent growth of NupFG condensates into larger structures. In line with this, our FRAP experiments of co-localizing Nup153FG and DNAJB6 assemblies in human cells suggest that DNAJB6 promotes and maintains a stable gel-like environment where Nup153FG molecules display slower mobility within the assemblies and slower exchange with the solution. This is consistent with recent atomic force microscopy experiments showing that co-localizing FUS and DNAJB6 condensates initially display a higher elastic response compared to FUS condensates alone, but this remains unchanged up to 48 hours, while FUS-only condensates transition to a solid-like state^57^.

To our knowledge, the detection of large assemblies of endogenous DNAJB6 in human embryonic stem cells is the first demonstration that endogenous DNAJB6 forms organized cytoplasmic assemblies under physiological conditions. Previously published SEC-SAXS data of *in vitro* DNAJB6 oligomers reported a maximum dimension of ∼30 nm for assemblies containing 55 subunits ^64^. Since the DNAJB6 particles analyzed here via STED microscopy significantly exceed this dimension, we hypothesize that these cytoplasmic assemblies represent higher-order clusters of individual DNAJB6 oligomers.

We additionally show that the closely related B’ JDPs, DNAJB2 and DNAJB8, also exhibit the ability to interact with and delay FG-Nup aggregation, and that their anti-aggregation efficiencies vary depending on the specific NupFG fragments involved. Despite being all LARKS-containing proteins^10^, we note that the number of LARKS motifs encoded in the IDRs of DNAJB6 and DNAJB8 (19) is higher than that of DNAJB2 (9). Nevertheless, the presence of LARKS alone does not necessarily justify the capacity of a protein to establish long-lived kinetically arrested structures as shown in Figure 7. Future research might answer if this difference explains why DNAJB6 and DNAJB8 form homotypic assemblies which only partly colocalize with NupFGs, while DNAJB2 forms mixed DNAJB2-NupFG assemblies.

### Sequence-encoded stability and evolutionary tuning

The dense gel-like phase of DNAJB6 presents a challenge: the high local concentration of residues capable of multivalent interactions inherently risks self-aggregation, which is incompatible with the dynamic properties required for chaperoning substrates. How is this encoded in the amino acid sequence? Our computational approach provides mechanistic support for the idea that not only amino acid composition, but crucially the spatial distribution of aromatic residues and LARKS within the IDR, determine the stability and reversibility of DNAJB6 assemblies. The DNAJB6-IDR resembles the sticker-and-spacer distribution in prion-like domains (PLDs), which are enriched in polar residues and are often interspersed by aromatic residues^89^. It has been proposed that the strong interactions among aromatic residues (stickers) are weakened by the favorable solvation of polar residues (spacers)^89^. In line with this, it has been shown that the uniform distribution of aromatic residues along PLDs favors solubility and LLPS over aggregation^71^. In fact, a proteome-wide analysis revealed that aromatic residues within human PLDs enriched in aromatic residues (> 10%) tend to be patterned in a non-random, uniform manner^71^. Interestingly, we find that DNAJB6-IDR is highly conserved in mammals, as well as the IDRs of the closely related B’ class JDPs DNAJB2 and DNAJB8. The high degree of primary amino acid sequence conservation of the DNAJB6-IDR is a remarkable feature considering that IDRs are generally known to conserve amino acid composition rather than position^72–76^, exemplified here with P53, but demonstrated for several other IDPs^70,73,90–92^.

Based on our mutational analysis, we speculate that altering the order of amino acids or the composition of the IDR may impede substrate interactions in two possible manners: either by primarily affecting DNAJB6 self-association kinetics and stability (i.e. DNAJB6^SCR1-3^) and/or by removing substrate interaction sites (i.e. DNAJB6^18xS/T>A^ and DNAJB6^12xF>A^). The latter likely not only alters its substrate affinity, and thus its ability to engage in heterotypic interactions, but may simultaneously alter its self-association properties. In line, *in vitro* data have shown that the association-dissociation dynamics of DNAJB6 are key to its ability to suppress Aβ fibril formation^66^. Our mutational analysis further revealed that the stability of DNAJB6 is easily compromised when altering the IDR amino acid sequence (while keeping the composition) as the mutants DNAJB6^SCR3^ and in particular DNAJB6^rL1^ form 0.5% SDS insoluble aggregates. The density, interaction strength and distribution of LARKS motifs is drastically altered in DNAJB6^SCR3^. Even those DNAJB6 variants that we created that maintained both the amino acid composition and distribution of the wildtype IDR, are not all fully functional. Particularly the fast-aggregating polyQ reveals the functional consequences of the subtle changes in these mutants possibly because polyQ, like suggested for Aβ^66^, may be especially dependent on rapid and efficient release of monomeric DNAJB6 from the gel-like assemblies. In contrast, the more stable, gel-like behavior of the FG-Nups may tolerate minor changes in DNAJB6 dynamics as they rather depend on the interactions with DNAJB6 gel-like assemblies. This could also explain the much lower *in vitro* stoichiometry ratios required for DNAJB6 to prevent polyQ or Aβ aggregation (1:10-100)^46,47,93^ than required for preserving FG-Nup disorder (1:1)^43^. We note that while our focus has been on the surveillance activity of chaperones against FG-Nup aggregation, it is also possible that this surveillance is bidirectional, in which both chaperones and FG-Nups benefit from interacting with each other.

### Physiological relevance and broader implications

We previously showed a role for DNAJB6 in the surveillance of FG-Nups in the timeframe between synthesis and assembly into NPCs, and confirmed the interaction between endogenous DNAJB6 and FG-Nups in the nucleus and cytoplasm of cells (see ^43^ and co-submitted Musskopf et al). Based on the current work, we suggest that the strong self-association behavior of DNAJB6 may also be implicated in pathological contexts such as dystonia, in which the accumulation of DNAJB6 within herniations may contribute to the observed sequestration of protein quality control components, potentially contributing to cellular dysfunction^45^. In this context, we speculate that the strong capacity of DNAJB6 to form gel-like assemblies contributes to the irreversible recruitment of proteins into NE herniations.

Gel-like assemblies are not unique to DNAJB6; they are also a feature of other protective cellular assemblies such as stress granules and P granules^94–100^. Future research may reveal that such gels do not always represent a transition to a non-functional state, but may more generally represent a stable protective state, combining high substrate avidity with dynamic exchange, thus preventing irreversible protein sequestration.

Finally, our current and previous findings^43,101–104^ that certain chaperones evolved to maintain protein disorder conceptually changes the view on the function of chaperones and extends it beyond their role in supporting and maintaining polypeptides folding into ordered domains. Our multiscale simulations further suggest that these chaperone functions emerge from finely tuned intermolecular interaction networks encoded within conserved disordered regions, linking sequence chemical make-up directly to the material properties of its assemblies and surveillance activity.

## Methods

### Protein purification

The expression and purification of yeast and human FG-Nup fragments (yNup60FG_aa389-511_, yNup100FG_aa1-580_, yNup116FG_aa1-725_, yNup145FG_aa1-219_, hNup153_aa875–1475_) and J-domain proteins (DNAJB1, DNAJB2a, DNAJB6b, DNAJB8 and mutants DNAJB6^18xS/T>A^, DNAJB6^12xF>A^, DNAJB1^DNAJB6-S/T^, DNAJB6^rL1^, DNAJB6^rL2^, DNAJB6^SCR1^, DNAJB6^SCR2^, DNAJB6^SCR3^) were performed as described before ^43^. In short, the pSF350 expression plasmid, containing a His6-tag at the N-terminal end and a cysteine residue at the C-terminal end was used for expression of the listed FG-Nup fragments and full-length JDPs (**Table 1**). For cell lysis and protein purification, a 100 mM Tris-HCl, 2M guanidine-HCl, pH 8.0 buffer was used. For labelling, the FG-Nup fragments were incubated with fluorescein-5-maleimide (5MF, Thermo-Scientific, 62,245) and JDPs with Alexa fluor 594 C5-maleimide (AF594, Alexa fluor 594 C5-maleimide, Thermo-Scientific, A10256). Proteins were concentrated using Vivaspin Protein Concentrator spin columns (Vivaspin 10/30 kDa MWCO Polyethersulfone, Cytiva, 28–9322–47, 28–9322–48) and stored at −80 °C at a final concentration of 100 μM in 100 mM Tris-HCl 2M guanidine-HCl, pH 8.0 buffer, supplemented with 10% glycerol. To reduce the impact of the fluorescent labels on the protein behavior, the fluorescein-5-maleimide-labeled FG-Nup fragments and Alexa fluor 594 C5-maleimide-labelled JDPs were mixed with unlabeled proteins at the ratios indicated in **Table 2**. Please note we do not quantify the labeling efficiency hence the fraction of labeled protein might be an overestimation.

**Table 1.**
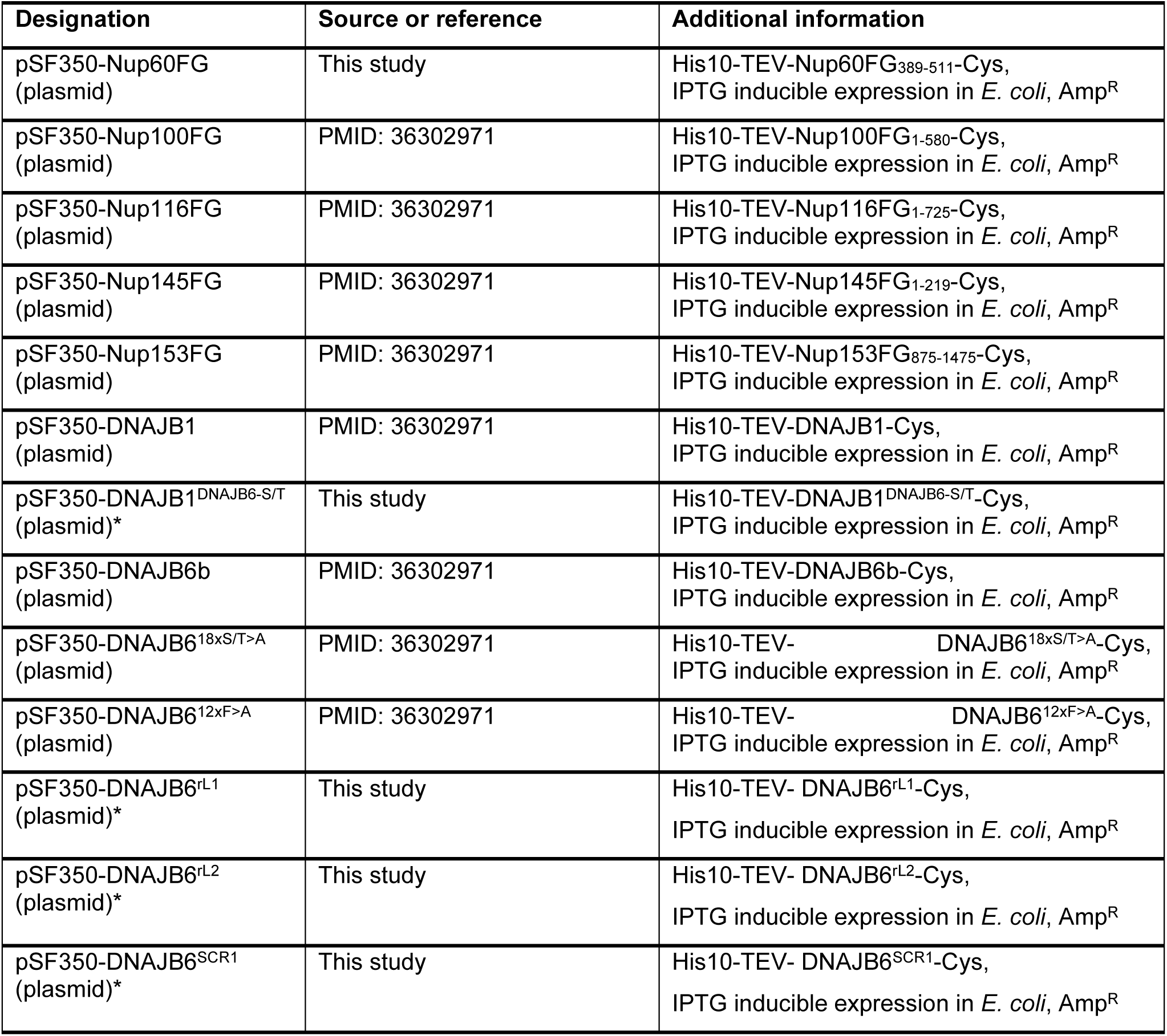
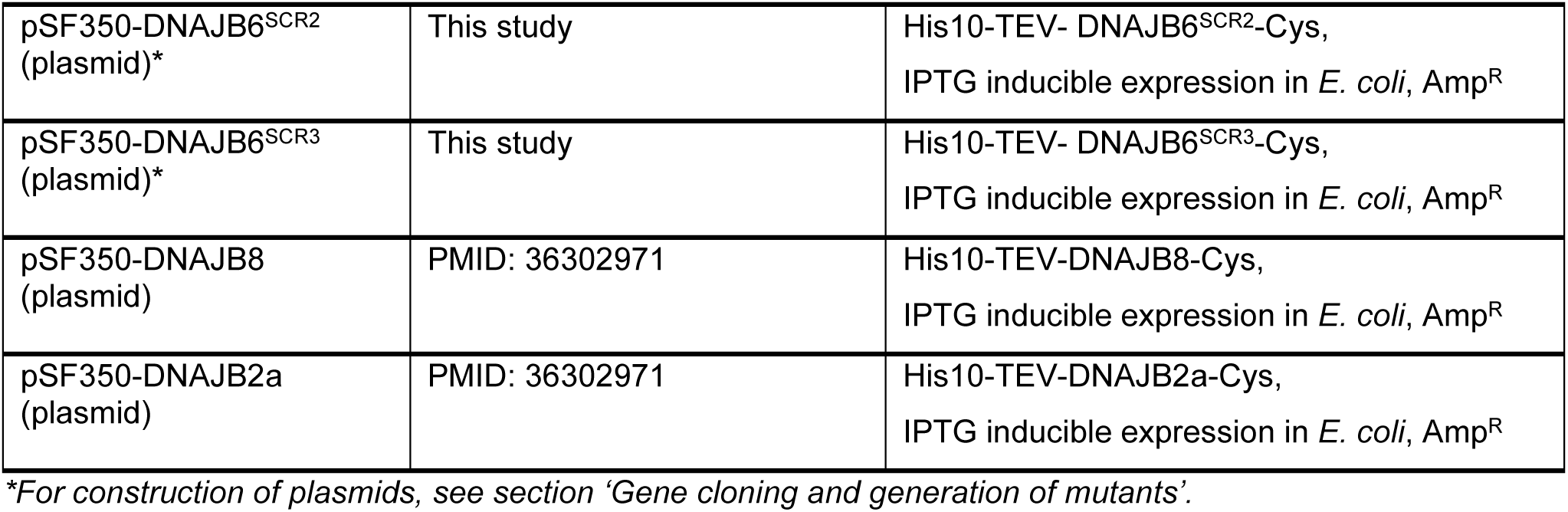
| Plasmids.

| Designation | Source or reference | Additional information |
| --- | --- | --- |
| pSF350-Nup60FG (plasmid) | This study | His10-TEV-Nup60FG<sub>389-511</sub>-Cys, IPTG inducible expression in *E. coli*, Amp<sup>R</sup> |
| pSF350-Nup100FG (plasmid) | PMID: 36302971 | His10-TEV-Nup100FG<sub>1-580</sub>-Cys, IPTG inducible expression in *E. coli*, Amp<sup>R</sup> |
| pSF350-Nup116FG (plasmid) | PMID: 36302971 | His10-TEV-Nup116FG<sub>1-725</sub>-Cys, IPTG inducible expression in *E. coli*, Amp<sup>R</sup> |
| pSF350-Nup145FG (plasmid) | PMID: 36302971 | His10-TEV-Nup145FG<sub>1-219</sub>-Cys, IPTG inducible expression in *E. coli*, Amp<sup>R</sup> |
| pSF350-Nup153FG (plasmid) | PMID: 36302971 | His10-TEV-Nup153FG<sub>875-1475</sub>-Cys, IPTG inducible expression in *E. coli*, Amp<sup>R</sup> |
| pSF350-DNAJB1 (plasmid) | PMID: 36302971 | His10-TEV-DNAJB1-Cys, IPTG inducible expression in *E. coli*, Amp<sup>R</sup> |
| pSF350-DNAJB1<sup>DNAJB6-S/T</sup> (plasmid)* | This study | His10-TEV-DNAJB1<sup>DNAJB6-S/T</sup>-Cys, IPTG inducible expression in *E. coli*, Amp<sup>R</sup> |
| pSF350-DNAJB6b (plasmid) | PMID: 36302971 | His10-TEV-DNAJB6b-Cys, IPTG inducible expression in *E. coli*, Amp<sup>R</sup> |
| pSF350-DNAJB6<sup>18xS/T>A</sup> (plasmid) | PMID: 36302971 | His10-TEV-DNAJB6<sup>18xS/T>A</sup>-Cys, IPTG inducible expression in *E. coli*, Amp<sup>R</sup> |
| pSF350-DNAJB6<sup>12xF>A</sup> (plasmid) | PMID: 36302971 | His10-TEV-DNAJB6<sup>12xF>A</sup>-Cys, IPTG inducible expression in *E. coli*, Amp<sup>R</sup> |
| pSF350-DNAJB6<sup>rL1</sup> (plasmid)* | This study | His10-TEV-DNAJB6<sup>rL1</sup>-Cys, IPTG inducible expression in *E. coli*, Amp<sup>R</sup> |
| pSF350-DNAJB6<sup>rL2</sup> (plasmid)* | This study | His10-TEV-DNAJB6<sup>rL2</sup>-Cys, IPTG inducible expression in *E. coli*, Amp<sup>R</sup> |
| pSF350-DNAJB6<sup>SCR1</sup> (plasmid)* | This study | His10-TEV-DNAJB6<sup>SCR1</sup>-Cys, IPTG inducible expression in *E. coli*, Amp<sup>R</sup> |
| pSF350-DNAJB6<sup>SCR2</sup><br>(plasmid)* | This study | His10-TEV- DNAJB6<sup>SCR2</sup>-Cys,<br>IPTG inducible expression in *E. coli*, Amp<sup>R</sup> |
| pSF350-DNAJB6<sup>SCR3</sup><br>(plasmid)* | This study | His10-TEV- DNAJB6<sup>SCR3</sup>-Cys,<br>IPTG inducible expression in *E. coli*, Amp<sup>R</sup> |
| pSF350-DNAJB8<br>(plasmid) | PMID: 36302971 | His10-TEV-DNAJB8-Cys,<br>IPTG inducible expression in *E. coli*, Amp<sup>R</sup> |
| pSF350-DNAJB2a<br>(plasmid) | PMID: 36302971 | His10-TEV-DNAJB2a-Cys,<br>IPTG inducible expression in *E. coli*, Amp<sup>R</sup> |

\*For construction of plasmids, see section 'Gene cloning and generation of mutants'.

**Table 2.** | Fraction fluorescently labeled proteins.

| Protein fragment | Volume fraction of labeled and unlabeled protein stocks each at 100 μM (maximum labelling %) |
| --- | --- |
| Nup60FG-5MF, Nup145FG-5MF | 2:3 (40%) |
| Nup100FG-5MF, Nup153FG-5MF | 1:4 (20%) |
| Nup116FG-5MF, All JDPs and corresponding mutants | 1:19 (5%) |

### Sample preparation for phase separation assays

To assay phase separation behavior, the purified proteins were diluted to a protein concentration of either 3 μM or 6 μM in assay buffer in low-protein-binding tubes. The assay buffer comprises 50 mM Tris-HCl, 150mM NaCl, pH8.0, supplemented with either no crowding agent or 10% w/v of polyethylene glycol 3350 (PEG3350; Sigma-Aldrich, P4338-2 KG) for varying amounts of time, as indicated in figure legends, at room temperature.

To assess the effect of the different JDPs and corresponding mutants on FG-Nup particle properties, 3 μM of 5MF labeled FG-Nup fragments and AF594 labeled JDPs were mixed at a 1:1 ratio and left to phase separate for 1 h at room temperature, unless stated otherwise in the figure legends. To assess the 1,6-hexanediol (HD) and SDS sensitivity of the different JDPs, particles were left to phase separate for 1 h, followed by exposure to either 5% 1,6-HD or 0.5% SDS for 10 min. For the Nup100FG:DNAJB6 time course experiment, Nup100FG particle formation was followed over a three-and-a-half-hour period, with 30 min time-intervals, in assay buffer containing 10% PEG3350, either in the absence or upon addition of DNAJB6 after 1 h of particle formation. In the DNAJB6-treated condition, the images corresponding to the 1 h time point were acquired immediately after DNAJB6 addition. For the experiments where we either let the FG-Nup or DNAJB6 pre-form particles prior to mixing, 3 μM of 5MF labeled FG-Nup fragments (Nup100FG or Nup153FG) and AF594 labeled DNAJB6 were either mixed from the start (1:1 ratio), or independently left to phase separate for either 30 min or 1 h at room temperature prior to mixing. After mixing, the samples were left for an additional hour, prior to microscopy or FTA assessments.

### Fluorescence microscopy in in vitro assays

For imaging of *in vitro* purified proteins, 2 μl samples of phase separated protein mixtures were mounted on untreated glass-slides with coverslip. Microscopy was performed in a temperature-controlled environment at 20 °C using a DeltaVision Deconvolution Microscope (Cytiva), using an Olympus UPLS Apo 100× oil-immersion objective (NA 1.4) and InsightSSITM Solid State Illumination using the FITC 525/48 and A594 625/45 filter sets. Detection was done with a either a CoolSNAP HQ2 or EDGE sCMOS5.5 camera. Fluorescence images were acquired with 30 Z-slices of 0.2 μm. Images were deconvolved using softWoRx software (Cytiva) and processed using the open-source software FIJI/ImageJ.

### Filter trap assays

For filter trap assays (FTA), protein samples were prepared as explained above, after which samples were added to 180 μl of sample buffer supplemented with 0.5% SDS and mixed by vortex before loading onto the Bio-Dot apparatus (Bio-Rad). FTAs were performed as described before ^43^. After this, membranes were blocked with 2.5% BSA in PBS-T (0.1%), incubated overnight with mouse primary antibody anti-His (1:5000, monoclonal mouse Tetra-His antibody, Qiagen, #34670), washed three times with PBS-T, incubated for 1h with secondary anti-mouse m-IgGk BP-HRP protein (Santa Cruz, sc-516102) (1:2500) and washed three times with PBS-T. Chemiluminescence was detected using enhanced chemical luminescence (ECL) reagent using the Chemidoc imaging system (BioRad). For the quantification of the FTA, band intensities were measured using FIJI and expressed relative to the average intensity of the control.

### Cell lines, cell culture, and transient transfections

HEK293T cells (Thermo Fisher) and HEK293T^DNAJB6-/-^ cells (see ^49^ for details on cell line generation) were maintained according to standard protocols. The cells were cultured in DMEM (Gibco) supplemented with 10% FCS (Greiner Bio-One) and penicillin/streptomycin (Invitrogen). For transient transfections, cells were grown to 70–80% confluence in a 37°C incubator with 5% CO2 and seeded into 6-well plates or 8-well chambered cover glass (Cellvis; C8-1.5H-N). For microscopy experiments requiring 6-well plates, coverslips coated with 0.0001% poly-l-lysine (Sigma) were used as the cell-seeding surface.

Human embryonic stem cell line 9 (Hues9) were a kind gift from Mathilde Broekhuis (iPSC CRISPR Facility at the European Research Institute for the Biology of Ageing). Hues9 cells were cultured on Matrigel (Corning, 15535739)-covered surface in StemFlex medium (Thermo Fischer Scientific, A3349401) and grown at 37°C and 5% CO2. Cells were dissociated with 0.5 mM EDTA (Gibco, 15575020) in PBS and split every 4-5 days in medium supplemented with 10 ng/ml Y-27632 (StemCell technologies, 72308).

Used siRNAs, all from Dharmacon/Horizon: SMARTpool siGENOME non-targeting (D-001206-13-20), SMARTpool Accell DNAJB6 siRNA (E-013020-00-0005), ON-TARGETplus TOR1A siRNA (J-011023-05-0002), ON-TARGETplus TOR1B siRNA (J-014203-09-0002), ON-TARGETplus TOR2A siRNA (J-015292-09-0002), ON-TARGETplus TOR3A siRNA (J-017824-11-0002).

### Gene cloning and generation of mutants

Details on the design and construction of V5-DNAJB6^WT^ and V5-DNAJB6^18xS/T>A^ constructs are available in ^48^ and ^46^. The GFP-Nup153FG construct is described in ^43^, while the GFP-Q71 construct is described in ^49^. V5-DNAJB6^12xF>A^ construct was created via site-directed mutagenesis using V5-DNAJB6^WT^ as the template and several mutagenesis rounds with different primer pairs. V5-DNAJB6^SCR1^, V5-DNAJB6^SCR2^, and V5-DNAJB6^SCR3^ mutants were made by enzyme digestion of V5-DNAJB6^WT^ and insertion of gBlocks (IDT) with mutations in the DNAJB6 sequence (amino acids 105-188):

SCR1: TSFSFSFFFHGSFFFSGNNDFTGSRTSRFRFSGGTSTTSFFLESGFMEFSSGIGFRKGDGFPSGGSGDGSLDTAGFGPRSPSSG

SCR2: SGPMGFGFSLTGGSRFTSFRFLEGSFGSPFNFFHSGRGPSGFGFRRFSSFDDSTSGFSDGAGGSGTPRSEGDFFDFNFGTFSS

SCR3: SDGEDRDTPFSGFGFGRESFHGSGFFFPGTFNFPSFGSRTFTPRSFSFGDGASGLNSGSGGGGTSSGFSLSGDFRRFMFSFSF

The DNAJB6^rL1^ and DNAJB6^rL2^ mutants were generated using the RandSeq tool from ExPASy and ordered using GeneArt. Codon usage was optimized for the entire sequence in GeneArt. In DNAJB6^rL1^ the residues in the S/T-rich region were completely scrambled, and in DNAJB6^rL2^ the F and G residues were deliberately separated (amino acids 105-188).

rL1 (randomized sequence): GRDPFSFDFFEDPFEDFFGNRRGPRGSRSRTFGGGFLPGFSSSSGDFSFTSGTSGSSFFAMFGGTTSSHFSLGFGSNGSFGGF

rL2 (randomized sequence): GRDPFSFDFFEDPFEDFFGNRRGPRGSRSRGSGSGSTFFAFPFFTFDGGSTFFLSGGHSSGGSLFMGSSGSTGSGSSGTFNFF

DNAJB1^DNAJB6 S/T^ (DNAJB1 flanks; DNAJB6 S/T region): GFPMGMGGFTNVNFGRSRSAQEPAGTGSFFSAFSGFPSFGSGFSSFDTGFTSFGSLGHGGLTSFSSTSFGGSGMGNFRKKQDPPVTHDLRVSL

Gibson assembly was used to replace the DNAJB6^WT^ S/T region with the generated randomized sequences, and to insert the S/T-rich region from DNAJB6 immediately after the G/F-rich region of DNAJB1.

### Immunofluorescence

Cells were rinsed with PBS to remove residual medium and fixed for 20 minutes using 4% paraformaldehyde (PFA) in PBS at room temperature (RT). After two washes with PBS, cells were permeabilized with 0.5% Triton X-100 for 15 minutes at RT and washed again with PBS. Subsequently, cells were washed twice with PBS+ (PBS supplemented with 0.5% bovine serum albumin and 0.15% glycine) before incubation with 80–200μl of primary antibodies diluted in PBS+ for 1 hour at RT or overnight at 4°C. Following four washes with PBS+, cells were treated with Alexa488-, Alexa594-, or Alexa633-conjugated secondary antibodies (Thermo Fisher Scientific) for 1.5–2 hours at RT in PBS+. Nuclei were stained using Hoechst 33342 dye (Life Technologies) at a 1:2,000 dilution in PBS for 5–10 minutes at RT. Finally, cells were washed twice with PBS and mounted using 80% glycerol or Mowiol® 4-88 (Carlroth) for imaging. For cells seeded in 8-well chambered cover glass, they were stored in PBS at 4°C until imaging. Images were acquired with Leica SP8 confocal microscope, using LAS-AFX software. Cells were imaged with 63x objective lens.

## STED Immunofluorescence

Cells were rinsed with PBS to remove residual medium and fixed for 20 minutes using 4% PFA in PBS at RT After three washes with PBS, cells were incubated in 100 mM glycine diluted in PBS for 10 minutes at RT. This was followed by two washes with PBS and cell permeabilization using 0.2% Triton X-100 for 15 minutes at RT. Cells were then washed once with PBS + 0.1% Tween (PBS-T) and blocked in 3% BSA diluted in PBS for 1 hour at RT. Primary antibodies were diluted in 3% BSA and 30 µL were added to the coverslip, which was covered with a small parafilm piece and incubated for 1 hour at RT. This was followed by three 5-minute washes with PBS-T. Secondary antibodies were also diluted in 3% BSA and 30 µL were added to the coverslip, which was covered with a small parafilm piece and incubated for 1 hour at RT, in the dark. Following the three 5-minute washes with PBS-T, nuclei were stained using Hoechst 33342 dye (Life Technologies) at a 1:2.000 dilution in PBS for 5 minutes at RT. Finally, cells were washed twice with PBS and mounted on 18 mm ∅ borosilicate glass coverslip, No. 1.5H using Mowiol® 4-88 (Carlroth). Cells were kept at RT overnight and then imaged.

STED images were acquired using a commercial STED microscope (Abberior Instruments GmbH), which has a 40 MHz pulsed 775 nm STED laser (3 W at laser head) and a pulsed 640 nm laser (1.2 mW at laser head) and is further equipped with a CoolLED pE-2 excitation system and a 100x oil immersion objective (Olympus UPLSAPO/1.40). STED images were taken using a pixel size of 15 nm, dwell time of 20 µs, 4 line repeats, 2% excitation power of the 640 nm laser and 40% STED depletion power. Image acquisition was carried out using Imspector Software (v16.3, Abberior Instruments).

### Image analysis

For the analysis of *in vitro* and in cells particle properties, the PhaseMetrics plugin^44^ in Fiji/ImageJ (version 1.54j) or CellProfiler (version 2.3.6) software was used. PhaseMetrics performs automatic detection and analysis of several particle properties, including particle intensity, size and circularity. A maximum intensity Z-projection is used for creating a segmented image for object detection, after which the object masks are re-directed to the sum of slices Z-projection to extract the desired measurements. To compute the overlapping area between FG-Nup and JDP particles, the object-based colocalization module was run to determine the intersected regions between the FG-Nup and JDP fluorescent signal. The accompanying “Colocalization analysis” excel macro was subsequently used to extract relevant colocalization-based measurements from the exported results table.

For the particle texture analysis, a custom CellProfiler pipeline was applied. The workflow included the following modules: *Smooth, Threshold, IdentifyPrimaryObjects, MeasureTexture, OverlayOutlines, SaveImages, and ExportToSpreadsheet*.

### FRAP analysis

HEK293T^WT^ cells were grown in 35mm poly-d-lysine coated glass bottom dishes (MatTek) before live cell imaging on a Zeiss780 confocal microscope in an incubation chamber at 37°C and 5% CO2. A FRAP region of interest covering a part of an accumulation of GFP-Nup153FG was selected and imaged for 10 iterations before the FRAP region was bleached for one iteration at the highest intensity of the 488 nm 25mW argon laser focused by a PlanApochromat 63x/1.40 Oil DIC lens. Recovery of fluorescence was monitored at the shortest interval possible (ms range) at 0.5% of the laser intensity used for bleaching. For generation of FRAP curves, the intensity values were fully normalized using the formula below

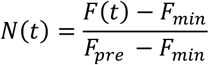

Where *N*(*t*) is the normalized intensity at time *t*, *F*(*t*) is the raw intensity at time *t*, *F_min_* is the intensity at the bleach frame, and *F_pre_* is the average intensity of the pre-bleach iteration.

### Antibodies

The primary antibodies used in this study, along with their respective companies, ordering numbers, host animals, western blot dilutions, and immunofluorescence dilutions, are listed below. mAb414 (Biolegend, 902901, mouse, -/1:500), GFP/YFP (Clontech, JL-8, anti-mouse, 1:5.000/1:500), MLF2 (Santa Cruz Biotechnology, sc-393566, mouse, -/1:500), GAPDH (Fitzgerald, 10R-G109A, mouse, 1:10.000/-), DNAJB6 (Braakman lab Utrecht, rabbit, 1:1.000/-), and V5 (Invitrogen, SY30-01, rabbit, 1:5.000/1:500). The following secondary antibodies and their respective company, ordering numbers, and dilutions were used. Anti-mouse-IgG HRP (GE Healthcare, GENXA931, 1:5.000), anti-rabbit-IgG HRP (GE Healthcare, GENA934, 1:5.000), Alexa Fluor 488 donkey anti-rabbit IgG (H + L) (Thermo Fisher, A21202, 1:500), Alexa Fluor 594 donkey anti-mouse IgG (H + L) (Thermo Fisher, A21203, 1:500), and Alexa Fluor 633 goat anti-rabbit IgG (H + L) (Thermo Fisher, A21070, 1:500), and for STED STAR 635 (Abberior, ST635, goat anti-mouse, 1:80).

### Protein fractionation

Cells grown in a 6-well plate were rinsed with PBS to remove residual medium and harvested in 500 µL PBS. After centrifugation at 500 g for 5 minutes at 4°C, the supernatant was discarded, and the cell pellet was lysed in 150 µL NP-40 lysis buffer (50 mM Tris-HCl pH 7.4, 100 mM NaCl, 1 mM MgCl2, 0.5% NP-40, 1x Protease Inhibitor, 50 U/mL DNArase, MilliQ). The lysates were vortexed thoroughly and incubated on ice for 30 minutes with periodic vortexing. Protein content was measured using the DC Protein Assay (Bio-Rad) and adjusted to a final concentration of 1–2 µg/µL. For fractionation, 15 µL of the lysate was taken as the whole-cell lysate (WCL). The remaining lysate was centrifuged at 14,000 g for 10 minutes at 4°C, and 15 µL of the supernatant was collected as the 0.5% NP-40 soluble fraction (S1). The leftover supernatant was discarded, and the pellet was washed by adding 150 µL Wash buffer (50 mM Tris-HCl pH 7.4, 100 mM NaCl, 1 mM MgCl2, MilliQ) and centrifuged at 14,000 g for 5 minutes at 4°C. The supernatant was discarded, and the pellet was resuspended in 100 µL of 0.1% SDS buffer (50 mM Tris-HCl pH 7.4, 100 mM NaCl, 1 mM MgCl2, 0.1% SDS, 1x Protease Inhibitor, MilliQ). This suspension was incubated in a ThermoMixer (Eppendorf) at 1,000 RPM and RT for 30–60 minutes, followed by centrifugation at 14,000 g for 10 minutes at RT. A 15 µL supernatant aliquot was collected as the 0.1% SDS soluble fraction (S2). The remaining supernatant was discarded, and the pellet was washed again with 150 µL Wash buffer and centrifuged as before. The pellet was then resuspended in 50 µL of 2% SDS buffer (50 mM Tris-HCl pH 7.4, 100 mM NaCl, 1 mM MgCl2, 2% SDS, 1x Protease Inhibitor, MilliQ) and incubated in the ThermoMixer at 1,000 RPM and RT for 30–60 minutes. After centrifugation at 14,000 g for 10 minutes at RT, a 30 µL supernatant aliquot was collected as the 2% SDS soluble fraction (P1). The pellet was washed again with 150 µL Wash buffer, centrifuged as before, and resuspended in 30 µL of 8M urea buffer to obtain the SDS-insoluble fraction (P2). This mixture was incubated overnight in the ThermoMixer at 1,000 RPM at RT. All aliquots (WCL, S1, S2, P1, and P2) were treated with 4X Laemmli buffer containing 10% β-mercaptoethanol, boiled at 99°C for 5 minutes, and loaded into TGX FastCast acrylamide gel (Bio-Rad, 1610172) for SDS-PAGE and western blot analysis.

### Evolutionary analysis

To assess the conservation of DNAJB6-IDR, two B’ JDP proteins (DNAJB2 and DNAJB8), and one unrelated IDP (P53) were used for a comparative analysis. A manually curated dataset of DNAJB6, DNAJB2, and DNAJB8 was utilized for subsequent steps, encompassing approximately 150 mammalian species. The UniProt IDs were uploaded to https://www.uniprot.org/peptide-search and separate files containing (i) length, (ii) region, (iii) organism (ID), (iv) sequence, (v) domain [FT], (vi) protein existence, and (vii) geneID were downloaded for DNAJB6, DNAJB2, and DNAJB8. A custom-made Python script was used to individually assess the Alphafold ^105^ files containing the per-residue pLDTT scores to identify the IDRs (pLDTT ≤ 65.0). For the JDPs, the end of the annotated J-domain (folded domain) served as a reference to identify their respective IDR1. Small, ordered regions (length ≤ 5) situated between IDRs were merged into the IDR, resulting in IDR1 sequences starting after the regulatory helix and ending just before the C-terminal β-sheet. Sequences without a predicted Alphafold structure, or with IDRs outside the length criterion (30 ≤ length ≤ 200), were excluded from the analysis. Subsequently, only species with an identified IDR1 for the three JDPs were retained (n=57). In these species, the C-terminal β-sheet was utilized as a reference to identify IDR2 from DNAJB6 (n=51) and DNAJB2 (n=57). A dataset for P53 in these 57 species was compiled manually, and IDR1 was identified using the beginning of the protein and the annotated DNA-binding domain (folded domain) as boundaries. Multiple sequence alignments (MSAs) of the whole protein, folded domain, IDR1, and IDR2 were generated using the default parameters of Clustal and visualized in the Jalview software. Conservation scores were calculated using the information_content ^106^ function from Bio.Align.AlignInfo module in Python, which returns a score per MSA column with a maximum number of 4.32 bits considering an equal probability for all 20 amino acids. A custom-made Python script was used to generate the IDR sequence logos using the logomaker package.

### Clustermap analysis of human IDRs

All human protein entries were retrieved from MobiDB databse (version 6.2; Release: 2024_07) ^107^. A custom Python script was used to analyze the corresponding AlphaFold structure files, extracting per-residue pLDDT scores to identify intrinsically disordered regions (IDRs), defined as contiguous segments with pLDDT ≤ 65.0. Short, ordered regions (≤ 5 residues) flanked by IDRs were merged to form continuous disordered segments. The NARDINI algorithm was applied to each identified IDR to compute z-scores for binary amino acid distribution and compositional fraction. Clustering of z-scores—either across the entire IDRome or within specific query sets—was performed using a custom Python script to generate clustermaps for comparative analysis.

### Statistical analysis

Graph preparation and statistical tests were performed using Prism10 software (GraphPad). The number of biological and technical replicates are indicated for all experiments in the figure legends. A color gradient was applied to highlight the individual datapoints belonging to each of the independent replicates. Data was tested for normality using the Shapiro-Wilk and D’Agostino-Pearson tests. For normal distributed data, a parametric unpaired t test (two groups) (**Figure 5N**) or one-way ANOVA (three or more groups) (**Figures S1G,H, Figure 3D**, **Figures 3P,Q, Figure S2F, Figures 4F-H**, **Figures 5F,G,I,J,L,M**) was performed. For non-normal distributed data, a nonparametric Mann Whitney test (two groups) (**Figure S4D**) or Kruskal-Wallis test (three or more groups) (**Figures 2B-D, Figures S1C,I-N, Figures 3I,J,K,M,N, Figures S2B,C,D,H,I,J, Figures S3C-H, Figure 6B, Figures S4A-C**) with Dunn’s multiple comparisons test was performed. Two-tailed p values; * p ≤ 0.05, ** p ≤ 0.01, *** p ≤ 0.001, ****p ≤ 0.0001.

### Molecular Dynamics simulations of DNAJB6 assemblies

Coarse-grained molecular dynamics simulations of DNAJB6 assemblies were performed using the residue-resolution Mpipi-Recharged force field, which represents each amino acid as a single interaction site. Hydrophobic interactions were modelled using the Wang-Frenkel potential with a cutoff of 3 s (i.e., molecular diameter of the beads; 1.5-1.7 nm), and electrostatic interactions among charged residues via a Yukawa potential (cut-off 3.5 nm), with a Debye screening length calculated self-consistently from the physiological salt concentration (150 mM NaCl) and the temperature-dependent relative dielectric permittivity. To account for buried interactions, the interaction energy between residues within globular structured domains was scaled by 0.7, and by the square root of 0.7 for pairs involving one structured and one disordered residue. Bonded interactions were modelled with a harmonic potential with force constant 9.6 kcal·mol⁻¹·Å⁻² and equilibrium bond length of 3.81 Å. All coarse-grained simulations were performed using the LAMMPS software ([LAMMPS](https://www.lammps.org/#gsc.tab=0)). The structured domains of DNAJB6 were treated as rigid bodies integrated with a Nosé-Hoover thermostat, while the disordered regions were propagated using a Langevin thermostat, both with a relaxation time of 5 ps and an integration timestep of 10 fs. The system pressure was controlled using a Berendsen barostat at isotropic pressure P = 0, with a relaxation time of 5 ps, consistent with the NpT approach used for critical temperature estimation of multicomponent condensates. Production runs were 1500 ns in timescale.

### Phase diagram

The phase diagram of DNAJB6 was determined from NpT bulk simulations at P = 0. Protein replicas were placed in a cubic simulation box and equilibrated across a range of temperatures. The critical temperature Tc was bracketed as lying between the highest temperature at which a stable condensed phase was observed and the lowest temperature at which the density diverged. The critical temperature and critical density were then extracted by fitting the law of rectilinear diameters and the law of critical exponents to the measured bulk densities, using the critical exponent α = 3.06 of the three-dimensional Ising universality class and assuming that the protein concentration in the dilute phase is negligible for our simulation box volumes.

### Intermolecular contact maps

Intermolecular contact maps were computed from simulation trajectories at temperatures of approximately 0.95*T_c_*, where *T_c_* corresponds to the critical temperature of each system. Contacts were identified using a sequence-dependent distance cut-off of 1.2σᵢⱼ, where σᵢⱼ represents the mean excluded volume of the respective i-th and j-th amino acids. This cutoff is set slightly beyond the potential minimum, located at approximately 2^1/6^*σ*ᵢⱼ ≈ 1.122*σ*ᵢⱼ, to ensure that only significantly bound pairs are counted.

### Atomistic potential of mean force calculations

LARKS segments were selected based on their predicted steric zipper propensity using ZipperDB ([ZipperDB](https://zipperdb.mbi.ucla.edu/)), retaining only sequences from the IDR sequence with an interface energy below −23 kcal·mol⁻¹ (see Appendix Figures S7D-H). Fibrillar assemblies for each selected LARKS were constructed using PyRosetta ([PyRosetta](https://www.pyrosetta.org/)) in a cross-β steric zipper configuration consistent with crystallographic templates, and pre-equilibrated for 3 ns at 300 K prior to free energy PMF calculations. All atomistic simulations using the Amber99sb-disp force field [https://www.pnas.org/cgi/doi/10.1073/pnas.1800690115] were performed using GROMACS 2023 ([gromacs](https://manual.gromacs.org/2023/)). Systems were solvated in cubic boxes of 7.5 × 7.5 × 7.5 nm at 150 mM NaCl, with ion parameters validated to reproduce aqueous solubility at 300 K. Energy minimization was performed to a maximum force below 1000 kJ·mol⁻¹·nm⁻¹, and all hydrogen bonds were constrained using the LINCS algorithm ([LINCS](https://onlinelibrary.wiley.com/doi/abs/10.1002/(SICI)1096-987X(199709)18:12%3C1463::AID-JCC4%3E3.0.CO;2-H)) with a 2 fs timestep. Long-range electrostatics were treated with Particle-Mesh Ewald ([PME](https://pubs.aip.org/aip/jcp/article-abstract/103/19/8577/180219/A-smooth-particle-mesh-Ewald-method)) using a real-space cut-off of 0.9 nm, and periodic boundary conditions were applied in all directions. Temperature and pressure were maintained with the velocity-rescale thermostat (*τ_T_* = 1 ps) and Parrinello–Rahman barostat (*τ_P_* = 1 ps), respectively. Potentials of mean force along the fibril separation coordinate were computed via umbrella sampling, with free energy profiles reconstructed using the weighted histogram analysis method ([WHAM](https://onlinelibrary.wiley.com/doi/abs/10.1002/jcc.540130812)). Total accumulated simulation time per system was 500 ns.

### Dynamic Algorithm for Ageing

Disorder-to-order structural transitions of DNAJB6 LARKS during condensate ageing were modelled using a time- and locally-dependent coarse-grained dynamic algorithm developed by us^77^. Every 100 simulation timesteps, the algorithm evaluates pairwise distances between LARKS of the same type at T = 280 K and triggers a disorder-to-order transition when two conditions are simultaneously satisfied: (1) two LARKS of the same type are within 11.0 Å of each other, and (2) both are surrounded by at least two further LARKS within 20.0 Å, with a cross-cutoff of 10.0 Å defining the local environment. Upon transition, LARKS residue identities are updated on-the-fly via the REACTION package of LAMMPS, with interaction energies, masses, and excluded volumes reassigned to values consistent with the structured β-sheet state evaluated through all-atom PMF simulations. To capture the accompanying increase in local rigidity, an angular harmonic term with equilibrium angle θ₀ = 180° and force constant k = 5 kcal·mol⁻¹·rad⁻² was applied to structured LARKS residues.

### Viscosity calculations

The shear viscosity was computed from the stress autocorrelation function evaluated at equilibrium bulk conditions corresponding to 0.95 T_c_. Simulations were performed in the NVT ensemble at the condensed-phase density obtained from the phase diagram, following compression of a Direct Coexistence box and relaxation in the NpT ensemble for ∼100 ns. Production runs for viscosity calculations ranged from 3 to 5 µs. The shear stress relaxation modulus G(t) was computed using six independent components of the pressure tensor via the multiple-tau correlator implemented in the EXTRA-FIX package of LAMMPS. The zero-shear viscosity η was obtained by integrating G(t) over time: at short timescales, numerical integration using the trapezoidal rule was applied; at longer timescales, where G(t) becomes noisy, the function was fitted to a series of Maxwell modes equidistant in logarithmic time using the open-source RepTate software ([RepTate](https://reptate.readthedocs.io/)), and the integral was evaluated analytically. The crossover time t₀ between the two regimes was chosen as the point beyond which all intramolecular oscillations in G(t) had decayed and the function became strictly positive and monotonically decreasing.

## Acknowledgements

We thank Johan Zijlstra for his initial characterization of the different FG-Nup fragments, Elizaveta Ustyantseva for providing Hues9 cells, and Tegan Otto for help with the STED imaging. We thank Michael Chang and all members of the Veenhoff, Kampinga, and Chang laboratories for their valuable suggestions. This work was financially supported by the Dutch Research Council (NWO) grant no. VI.C.192.031 to L.M.V. (L.M.V., A.S.) and OCENW.GROOT.2019.068 to H.H.K and L.M.V (T.B., M.M.K, P.G), project 95169 from the Dutch Campaign Team Huntington in the Netherlands and the NWW-funded NWA-ORC project “Cure-Q” to H.H.K, and a PhD fellowship from the Graduate School of Medical Sciences from the University Medical Center Groningen to M.M.K. We thank the Center for Information Technology of the University of Groningen for their support and for providing access to the Hábrók high performance computing cluster. We also gratefully acknowledge the Isambard supercomputing facility [https://docs.isambard.ac.uk/] and the MareNostrum 5 supercomputer at the Barcelona Supercomputing Center through RES computational resources (activities 2024-3-0001 and 2025-1-0009) for providing the computing time necessary for the molecular dynamics simulations. JRE acknowledge funding from the Ramón y Cajal fellowship (RYC2021-030937-I), the Spanish scientific plan and committee for research reference PID2022-136919NA-C33, and European Research Council (ERC) under the European Union’s Horizon Europe research and innovation programme (Grant Agreement No. 101160499).

## Author contributions

TB (conceptualization, formal analysis, investigation, methodology, visualization, writing original draft and review and editing); MKM (conceptualization, formal analysis, investigation, methodology, visualization, writing original draft and review and editing); AF (formal analysis, investigation, methodology, visualization review and editing); PG (formal analysis, investigation, methodology, visualization, review and editing); MER (data curation, review and editing); EFEK (formal analysis, investigation); NHE (formal analysis, investigation, methodology, visualization, review and editing); ART (formal analysis, investigation, methodology, visualization, review and editing); JF (formal analysis, investigation); SMYF (formal analysis, investigation); AS (supervision, review and editing); RV (formal analysis, methodology); JRE (conceptualization, supervision, funding acquisition, review and editing); HHK (conceptualization, supervision, funding acquisition, review and editing); LMV (conceptualization, supervision, funding acquisition, review and editing).

---

**Figure S1.**
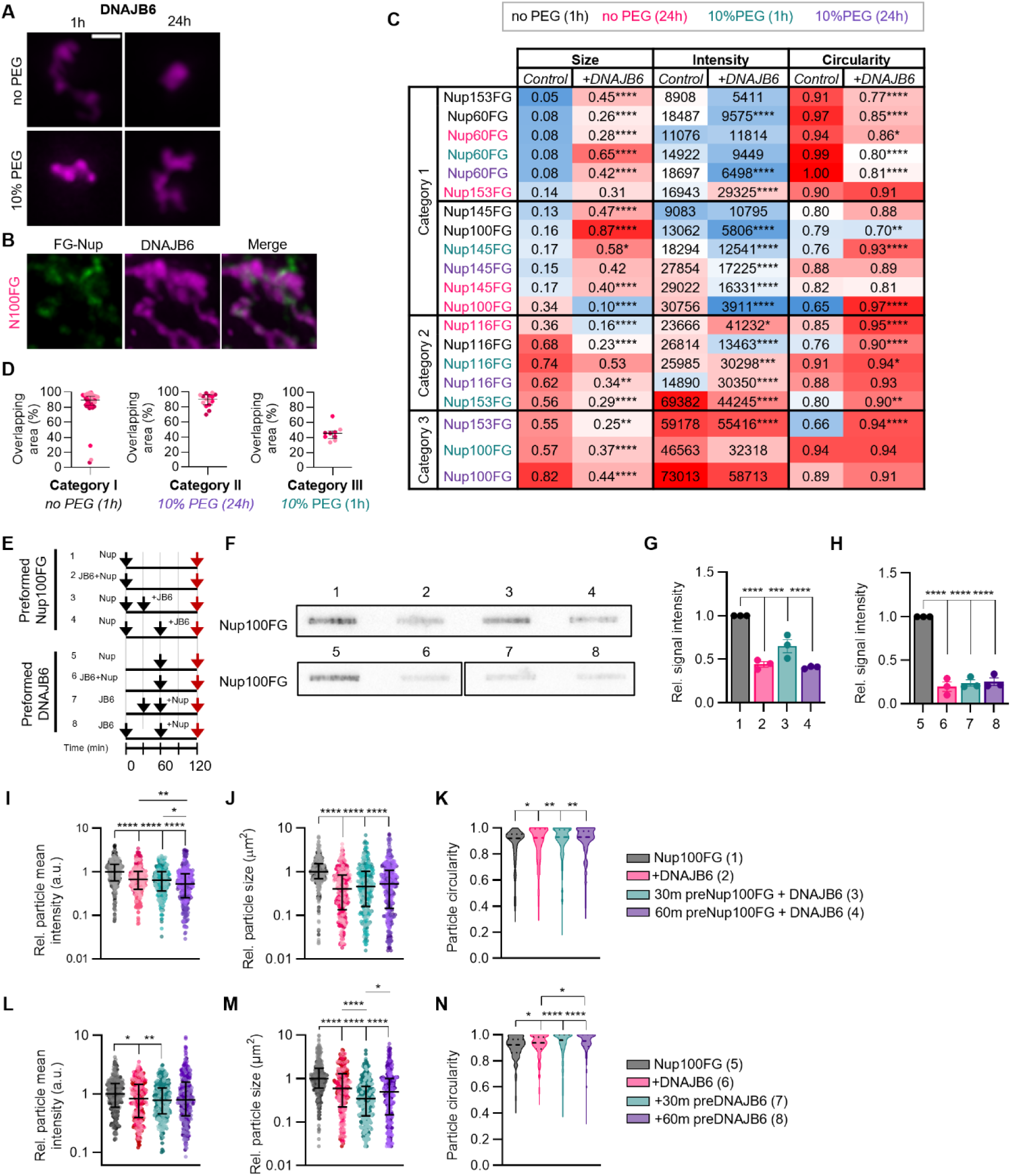
DNAJB6 can modulate a range of NupFG condensates. **(A)** Representative images showing DNAJB6-A594 particles, formed in the absence or presence of 10% PEG3350 for 1h or 24h. Scale bar, 1 µm. **(B)** Few DNAJB6 particles were observed in the Nup100FG-5MF:DNAJB6-A594 mixture formed in the absence of 10% PEG3350 after 24h (as visualized in Figure 2A), but in rare cases DNAJB6 particles remained and interacted with Nup100FG as exemplified in the image. **(C)** Table showing overview of median particle size, intensity and circularity for each of the conditions exemplified in (Figure 2A), sorted in the same order. A colour-gradient was applied to each of the assessed particle properties, with lower values indicated in blue and higher values indicated in red. ≥300 particles per condition (n=4). Stars indicate significance. **(D)** Percentage of overlapping areas between DNAJB6-A594 and NupFG-5MF particles represented in (Figures 1E-G). **(E)** Schematic diagram of mixing conditions for experiments depicted in (F-H). Red arrow indicates time of imaging. **(F)** Filter trap assay showing aggregated fraction of Nup100FG in the absence or presence of DNAJB6 (molar ratio 1:1). Numbers refer to conditions in (E). **(G,H)** Quantification of the band intensities of Nup100FG on filter trap for each of the conditions exemplified in (F). Represented band intensities are relative to the average intensity of the control. Mean ± SEM (n=3). **(I-N)** Mean fluorescence intensity (I,L), size (J,M) and circularity (K,N) of Nup100FG-5MF particles exemplified in (Figure 1M), in which proteins were either mixed from the start or either Nup100FG (top) or DNAJB6 (bottom) was pre-assembled for either 30 or 60 min prior to mixing (molar ratio 1:1). The mean fluorescence intensity and size data was plotted relative to the median of the control. Graphs show median ± interquartile range of ≥300 particles per condition (n=3). *P<0.05, **P<0.01, ***P<0.001, ****P<0.0001.***P<0.001, ****P<0.0001.

**Figure S2.**
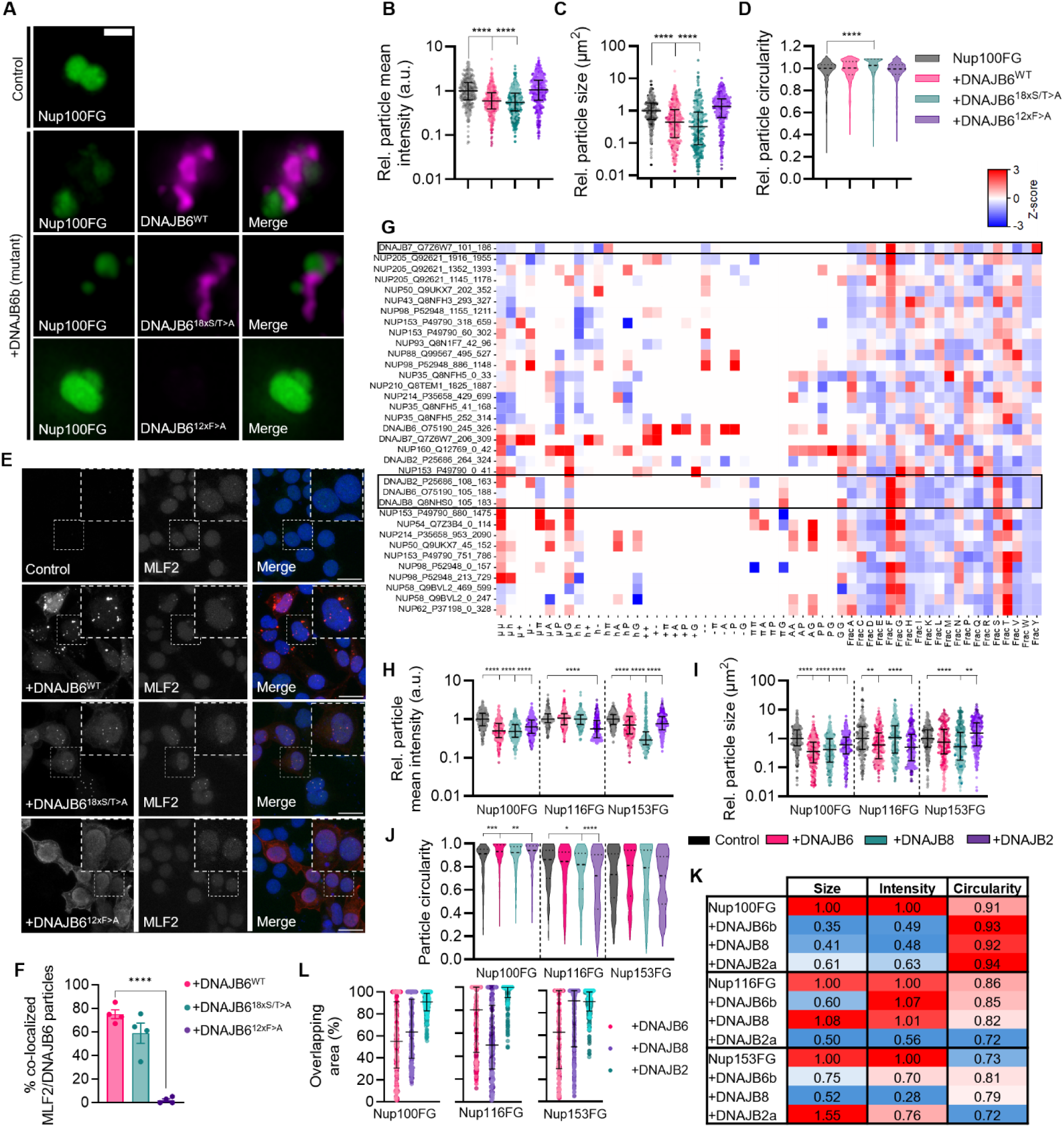
Sequence-encoded ability to regulate FG-Nup phase transitions. **(A)** Representative images showing Nup100FG-5MF particles in the absence or presence of either DNAJB6^WT^, DNAJB6^18xS/T>A^ or DNAJB6^12xF>A^ particles, formed in the presence of 10% PEG3350 (1h) (molar ratio 1:1). Scale bar, 1 µm. **(B-D)** Mean fluorescence intensity (B), size (C) and circularity (D) (relative to the median of the control) of Nup100FG-5MF particles exemplified in (A). Graphs show median ± interquartile range of ≥300 particles per condition (n=4). **(E)** Representative images of HEK293T^WT^ cells transfected with V5-tagged DNAJB6b constructs (DNAJB6^WT^, DNAJB6^18xS/T>A^, or DNAJB6^12xF>A^). Scale bar, 20 µm. **(F)** Quantification of MLF2 foci co-localized with DNAJB6 foci exemplified in (E). Graphs show mean ± SEM (n=4). **(G)** Clustermap of z-scores from NARDINI algorithm for binary amino acid distributions and amino acid fractions in the human IDRs of DNAJB2, DNAJB6, DNAJB7, DNAJB8, and FG-Nups. Rectangles indicate the IDRs of B’ JDPs located between the regulatory helix and the C-terminal β-sheets. **(H-J)** Mean fluorescence intensity (H), size (I) and circularity (J) of Nup100FG-5MF, Nup116FG-5MF and Nup153FG-5MF particles exemplified in (Figures 4I-K). Graphs show median ± interquartile range of 300 particles per condition (n=3). The mean fluorescence intensity and size data was plotted relative to the median of the control. **(K)** Table showing overview of median particle size, intensity and circularity for each of the conditions exemplified in (Figures 4I-K). A colour-gradient was applied to each of the assessed particle properties, representing the magnitude of change relative to the control condition, with lower values indicated in blue and higher values indicated in red. **(L)** Graph showing the median overlapping area for Nup100FG, Nup116FG, and Nup153FG with DNAJB6, DNAJB8, and DNAJB2 conditions exemplified in (Figures 4I-K). *P<0.05, **P<0.01, ***P<0.001, ****P<0.0001.

**Figure S3.**
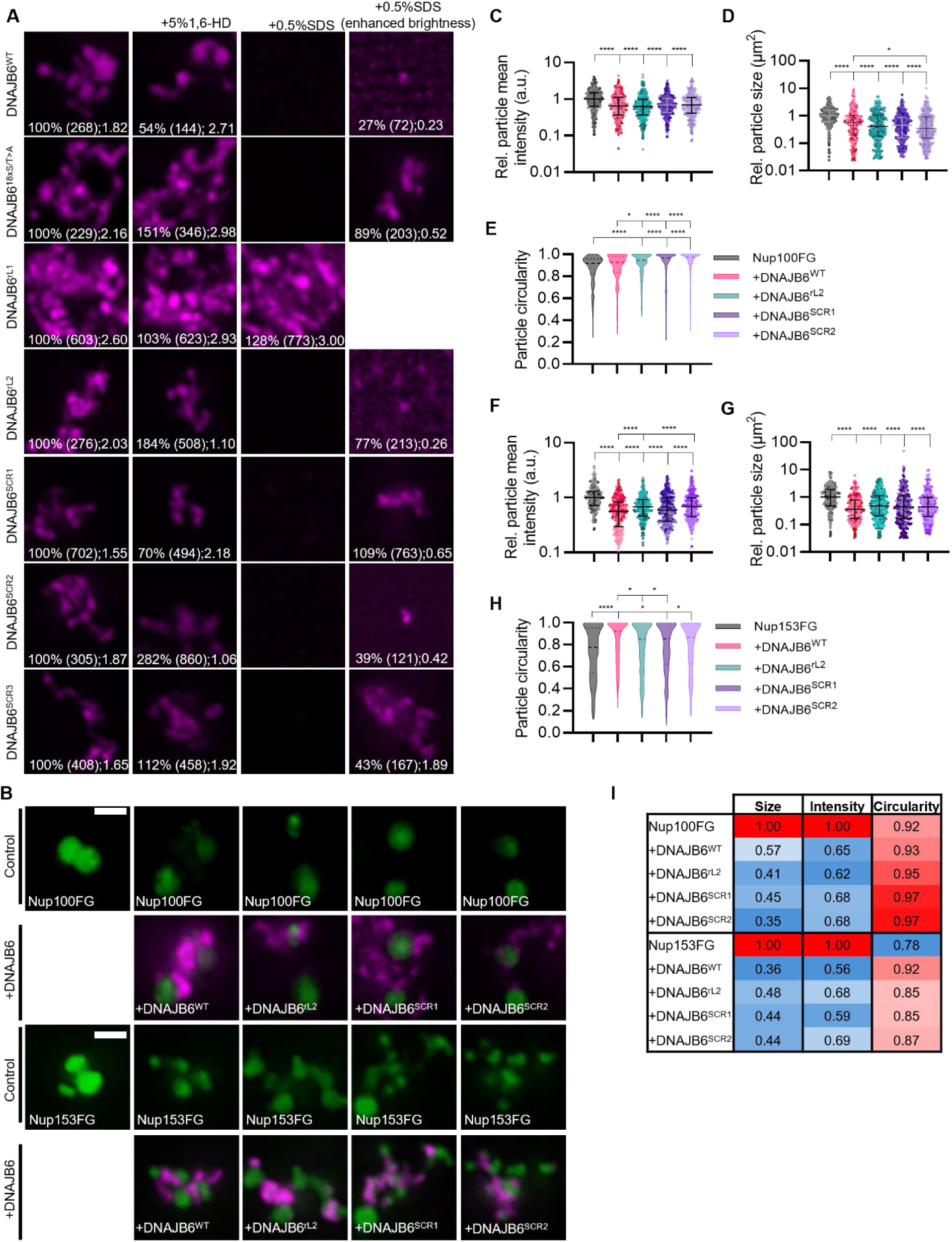
Alterations in the DNAJB6-IDR sequence impact DNAJB6 its ability to form stable assemblies. **(A)** Representative images of *in vitro* purified DNAJB6^WT^ and DNAJB6^rL1^, DNAJB6^rL2^, DNAJB6^SCR1^, DNAJB6^SCR2^ and DNAJB6^SCR3^ particles, formed in the presence of 10% PEG3350 (1h). After 1 hour incubation, samples were treated with either 5% 1,6-hexanediol or 0.5% SDS for 10 minutes. Numbers indicate the percentage of particles remaining after chemical treatment relative to the control sample, total number of particles counted and median particle size (n=2). **(B)** Representative images showing Nup100FG-5MF and Nup153FG-5MF particles in the absence or presence of either DNAJB6^WT^, DNAJB6^rL2^, DNAJB6^SCR1^ or DNAJB6^SCR2^ (1h) (molar ratio 1:1). Scale bar, 1 µm. **(C-H)** Mean fluorescence intensity, size and circularity of Nup100FG-5MF (C-E) and Nup153FG-5MF (F-H) particles exemplified in (B). The mean fluorescence intensity and size data was plotted relative to the median of the control. Graphs show median ± interquartile range of 300 particles per condition (n=3)). **(I)** Table showing overview of median particle size, intensity and circularity for each of the conditions exemplified in (B) and quantified in (C-H). A colour-gradient was applied to each of the assessed particle properties, representing the magnitude of change relative to the control condition, with lower values indicated in blue and higher values indicated in red. *P<0.05, **P<0.01, ***P<0.001, ****P<0.0001.

**Figure S4.**
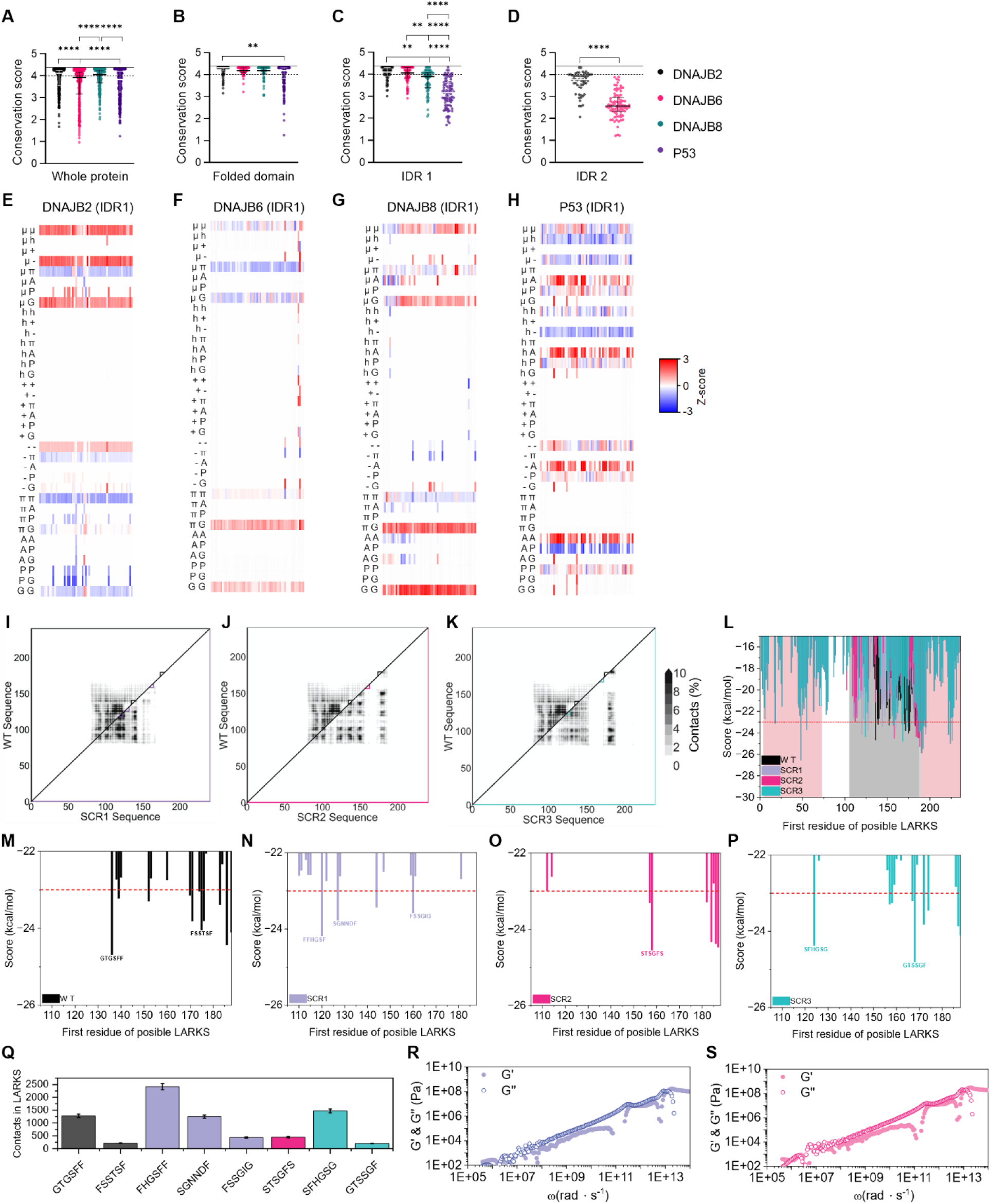
Evolutionary and coarse-grained analyses. (**A-D**) Conservation scores of residues from the whole protein (A), folded domain (B), IDR1 (C), or IDR2 (D) from DNAJB2, DNAJB6, DNAJB8, and P53. Graphs shows median with interquartile range. **P<0.01, ***P<0.001, ****P<0.0001. (**E-H**) Z-scores for the binary amino acid distributions in the IDR1 of DNAJB2 (E), DNAJB6 (F), DNAJB8 (G), and P53 (H) across the 57 analyzed mammal species. **(I-K)** Intermolecular contact frequency maps comparing the wild-type (WT) sequence against each scrambled variant: SCR1 (I), SCR2 (J), and SCR3 (K). The upper triangle displays WT contact frequencies and the lower triangle displays the corresponding scrambled variant contact frequencies, allowing direct visual comparison of inter-residue interaction patterns. **(L)** ZipperDB aggregation scores as a function of the first residue of each candidate LARKS hexapeptide across the full protein sequence for WT (black), SCR1 (purple), SCR2 (pink), and SCR3 (cyan). Grey and red background regions denote the IDR and structured segments, respectively. The red dashed line indicates the ZipperDB score threshold (-23 kcal/mol) below which hexapeptides are considered aggregation-prone. **(M-P)** ZipperDB score profiles restricted to the IDR region for WT (M), SCR1 (N), SCR2 (O), and SCR3 (P), with the most aggregation-prone LARKS sequences labeled explicitly. **(Q)** Total number of intermolecular contacts formed within the predicted LARKS regions for each identified hexapeptide across all variants, showing the relative propensity of each segment to engage in fibril-like interactions. Error bars represent standard deviation across simulation replicates. **(R,S)** Viscoelastic moduli G′ and G″ as a function of frequency for SCR1 (R) and SCR2 (S) variants, derived from the stress tensor.

